# Decoupling between activation time and steady-state level in input-output responses

**DOI:** 10.1101/2025.09.03.673941

**Authors:** Giorgio Ravanelli, Kee-Myoung Nam, Jeremy Gunawardena, Rosa Martinez-Corral

## Abstract

Many biological processes, like gene regulation or cell signalling, rely on molecules (inputs) that bind to targets leading to downstream responses. In the gene regulation field, recent data have shown that higher transcription factor (TF) concentrations may increase transcription levels of a gene without affecting the gene activation time. We call this behaviour *output decoupling*. Motivated by these observations, here we investigate mechanisms for output decoupling in Markov process models where a readout molecule is produced downstream of ligand binding. Our focus is on identifying regimes where the steady-state level of the readout changes with input concentration, while the activation time, quantified by mean first-passage times, remains unaffected. Through a combination of analytical and numerical investigations, we find two mechanisms through which output decoupling can arise: i) *rate scale separation*, where the system is comprised of slow and fast transitions that are differentially regulated by the input; and ii) incoherent regulation, where the input acts on two transitions with opposing effects on readout production, with all transitions operating on similar timescales in the absence of input. Such incoherent regulation has emerged as a plausible regulatory mode of TFs, and we suggest decoupling as a new characteristic feature of this regulatory mode. More broadly, our findings offer a mechanistic and conceptual framework for reasoning about output decoupling in input-output systems.

**Author summary:** How biomolecular systems respond to signals often depends not only on the final level of activity they reach, but also on how quickly they reach it. These two outputs — steady-state level and activation time— are usually coupled: higher input concentration tends to produce both higher activity and faster responses. Yet recent experiments suggest that, in some cases, the strength of the response can change while the speed remains constant. In this work, we explore the conditions under which such “output decoupling” can occur. Using mathematical models of molecular systems, we identify two ways this can happen. In one case, the activation time is determined by slow steps in the system that the input does not control. In the other, the input exerts opposing effects on the system, simultaneously promoting and hindering the response, which balances the timing. By revealing how decoupling arises, our study provides a framework for interpreting puzzling experimental results. More broadly, it shows the value of considering both dynamics and steady-state behavior jointly when studying how molecular systems process information.

## Introduction

Many biological processes are regulated by an input molecule that binds to a target. Examples include ligands binding to receptors, transcription factors binding to regulatory sites on DNA, and splicing regulators binding to pre-mRNAs. Upon binding, the molecular system can undergo internal molecular changes that result in a measurable readout (Fig. 1A). Example readouts can be a receptor’s phosphorylation level, a gene’s expression level, or an exon’s inclusion level. Usually, we extract summary features, or “outputs”, from the molecular readout and study their relationships with the input levels (Fig. 1A,B,C). We call these types of mapping input-output responses [1–5].

**Fig. 1.**
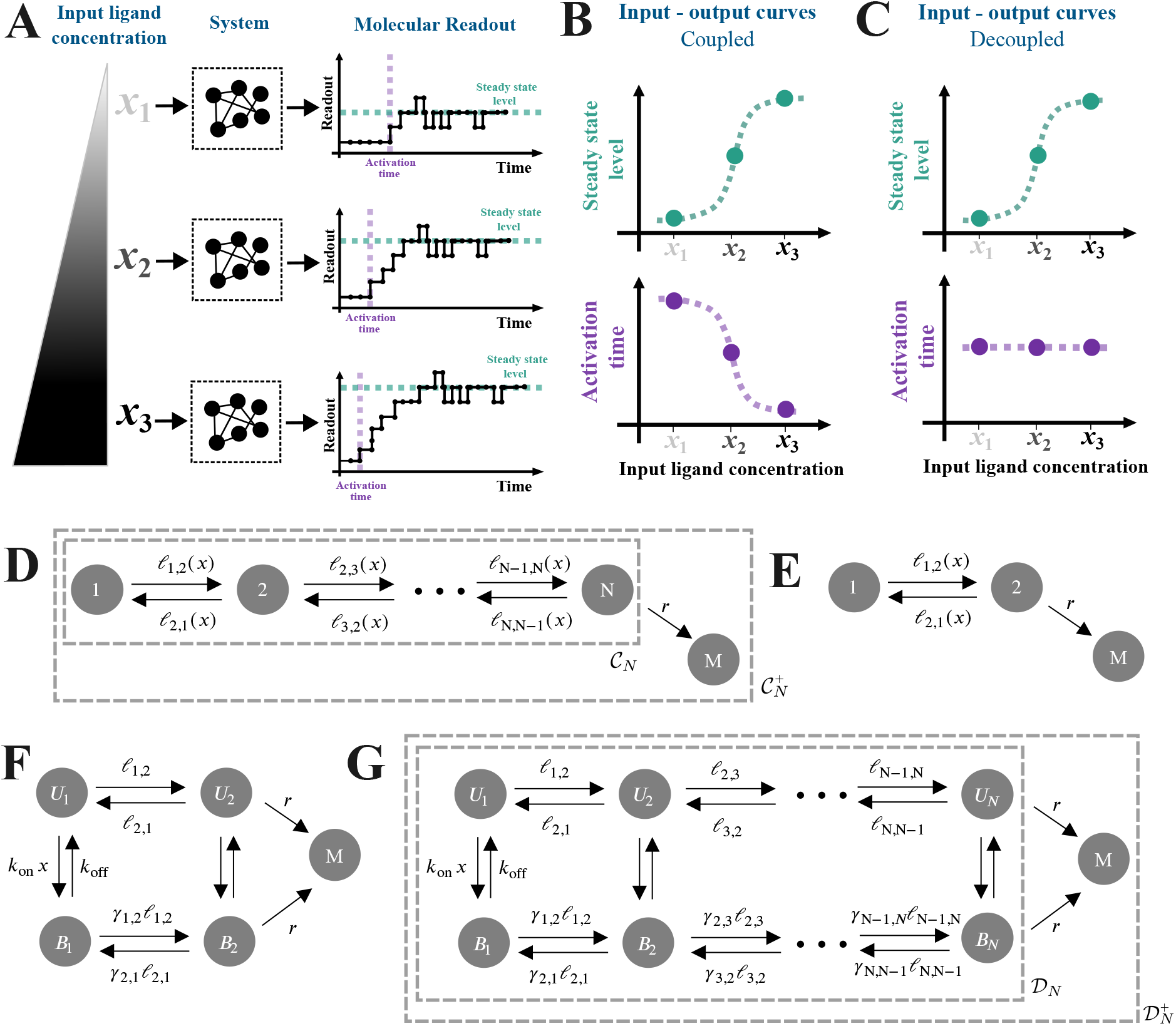
Interplay between steady state and activation time. **(A)** Schematic of input-output responses. An input ligand with concentration *x* (left) is processed by a system from which we measure a molecular readout, from which the readout’s steady-state level or activation time can be quantified (right). Here, “activation time” is defined as the time required for the readout to increase by one molecule after the input has been introduced. **(B–C)** Schematic of coupled (**B**) and decoupled (**C**) input-output responses for the steady-state level and activation time. In the former, the steady-state level increases while the activation time decreases with input concentration; in the latter, the steady-state level increases with the input concentration, while the activation time remains constant. **(D–G)** The models used in this paper, where *M* is the molecular readout and *x* denotes the input concentration. See text for more details. **(D–E)** Chain models with implicit ligand binding. The ligand’s regulatory effect is captured in the edge labels as arbitrary functions of *x*. The graph, 𝒞_*N*_, without the terminal state *M* is used to calculate the steady-state level; the augmented graph, 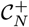, is used to calculate the activation time. See text for more details. **(F–G)** Ladder models that explicitly incorporate ligand binding. The vertical edges represent ligand binding and unbinding, with rates *k*_on_*x* and *k*_off_, respectively. As with the chain models, the graph, 𝒟_*N*_, without the terminal state *M* is used to calculate the steady-state level, while the augmented graph, 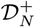, is used to calculate the activation time. See text for more details.

Two outputs commonly considered in the literature are the *steady-state* level of the molecular readout, and the *activation time*. The latter can be defined and quantified in different ways (Discussion). In this work, we consider the activation time as the time for the molecular readout to change upon introducing the input (Fig. 1A), and we will define it formally below. When both the steady-state and activation time have been quantified as functions of input concentration, experiments have typically shown *coupling* : a higher steady-state level is usually accompanied by faster (smaller) activation times (Fig. 1B). Examples include *β*-adrenergic receptor activity as a function of drug concentration [6], viral entry as a function of receptor concentration [7], and transcription of various genes as a function of TF concentration [8–13].

Coupling can be intuitively explained by thinking about a system that undergoes a series of reversible transitions between an inactive and an active state from which the molecular readout is produced, with the binding of the input facilitating (accelerating) the transitions towards the productive state [9, 14, 15]. In this scenario, higher input concentrations reduce the time for the readout to change while also increasing its steady-state level.

By contrast, experimental data from recent studies of eukaryotic gene regulation suggest the possibility of *output decoupling*, where the steady-state transcription level of a gene increases as a function of the concentration of an input TF, but the activation time does not change [9, 11, 14, 16] (Fig. 1C). There are some subtleties in these data and how they are analysed, which we will turn to in the Discussion. Leaving these aside for now, the general question remains: what kinds of regulatory mechanisms can underlie output decoupling?

To explore this question, we focus on processes in which a ligand binds to a target and promotes the downstream production or accumulation of a molecule. This accounts for TFs activating transcription, or a ligand-bound receptor influencing protein modification or altering the permeability of a channel or transporter. For generality, we use the term “ligand” to refer to the input molecule, although we mostly frame the work in the language of gene regulation, given the motivating observations in that field.

We begin by giving a general overview of the models, namely Markov processes, that we examine and the mathematical formalism, namely the graph-theoretic *linear framework* [17–20], that we use to analyse them. We start with the simple two-state telegraph model, then proceed to more complex models. By employing a dialogue between numerical approaches and analytical calculations, we find two regulatory strategies for output decoupling: i) when the system exhibits *rate scale separation*, in which a slower transition or set of forward transitions govern the activation time, with the input affecting the steady state by modulating other transitions and ii) when the system exhibits an *incoherent regulatory mode*, which simultaneously promotes and hinders production of the readout. This regulatory mode has recently garnered attention in the field of eukaryotic gene regulation [21–23], and our results suggest that output decoupling could be another significant consequence of such incoherent regulation. More generally, we demonstrate that a rich mechanistic playground is uncovered when jointly investigating the steady state and the transient regime of molecular systems, thus further motivating joint experimental and theoretical investigation of these two regimes.

### Modeling approach and mathematical setup

We model ligand-binding-readout systems as discrete-state, continuous-time Markov processes using the graph-theoretic linear framework [17–20]. Specifically, we describe the system as a finite, directed graph, *G*, with labelled edges, in which the vertices, 𝒱 (*G*), represent the system states; the edges, denoted *i* → *j*, represent transitions among the states; and the edge labels, denoted *ℓ*(*i* →*j*), represent transition rates with dimensions of (time)^*−*1^. Such a description gives rise to a corresponding continuous-time Markov process, *X*(*·*), on state space 𝒱 (*G*), in which the infinitesimal transition rate from *i* to *j* is nonzero if, and only if, the edge *i* →*j* exists in *G*, and the rate is given by the edge label:

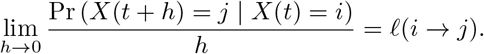

Now, suppose that 𝒱 (*G*) = {1, …, *n*}, and let *p*_*i*_(*t*) be the probability that *X*(*·*) occupies vertex *i* ∈ 𝒱 (*G*) at time *t*, given some choice of initial vertex. The time-evolution of this probability is given by the master equation [18, 24],

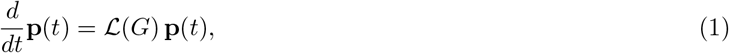

where **p**(*t*) = (*p*_1_(*t*), …, *p*_*n*_(*t*))^T^ and ℒ (*G*) is the *n × n Laplacian matrix* of *G*, whose entries are given by

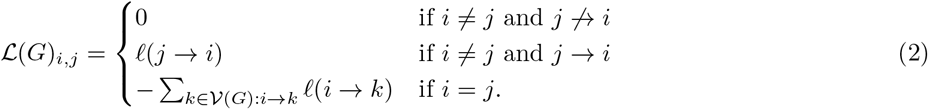

At steady state, the left-hand time derivative in Eqn. 1 can be set to zero. Therefore, the vector of steady-state probabilities, which we denote by **p**^*∗*^, lies in the kernel of ℒ (*G*). We shall extensively refer to this master equation in what follows.

We consider two kinds of models in this paper. First, we consider *chain models* (also called *pipeline models* in previous work [20]), denoted 𝒞_*N*_, in which *N* vertices are reversibly connected in a linear arrangement. In particular, the vertices of *C*_*N*_ are given by 𝒱 (𝒞_*N*_) = {1, …, *N*}, and consecutive vertices are connected by reversible edges, *i → i* + 1 and *i* + 1 *→ i* for *i* = 1, …, *N −* 1 (Fig. 1D–E). We introduce the ligand implicitly through the edge labels, denoted *ℓ*(*i* →*j*) = *ℓ*_*i,j*_(*x*), which depend on the ligand concentration, *x* (Fig. 1D–E) [25]. With this parametrization, many possible functional forms for the dependence on *x* are biologically plausible.

In order to formalise a particular assumption for how the ligand influences the system, we then turn to *ladder models*, denoted 𝒟_*N*_, which explicitly incorporate ligand binding and assume that the ligand has an effect while bound (Fig. 1F–G) [13, 26]. In particular, 𝒟_*N*_ is a graph on 2*N* vertices,

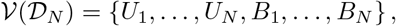

where *U*_1_, …, *U*_*N*_ represent ligand-unbound states and *B*_1_, …, *B*_*N*_ represent ligand-bound states. There are edges between pairs of consecutive unbound vertices, *U*_*i*_ *→ U*_*i*+1_ and *U*_*i*+1_ *→ U*_*i*_; pairs of consecutive bound vertices, *B*_*i*_ *→ B*_*i*+1_ and *B*_*i*+1_ *→ B*_*i*_; and pairs of unbound and bound vertices of the same index, *U*_*i*_ *→ B*_*i*_ and *B*_*i*_ *→ U*_*i*_. We assume that the label on each binding edge, *U*_*i*_ *→ B*_*i*_, and unbinding edge, *B*_*i*_ *→ U*_*i*_, is independent of the index *i*, as

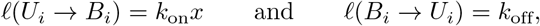

where *x* is the ligand concentration, *k*_on_ is a rate constant with units of (concentration · time)^*−*1^, and *k*_off_ is an off-rate. On the other hand, we allow for the other edge labels to vary with *i*, and write them as

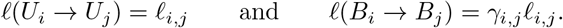

As such, *γ*_*i,j*_ *>* 0 is a dimensionless parameter that captures the extent to which the ligand promotes (*γ*_*i,j*_ *>* 1), hinders (*γ*_*i,j*_ *<* 1), or maintains as is (*γ*_*i,j*_ = 1) the transition from index *i* to index *j*. We call these parameters regulatory factors.

Each system we consider in this paper produces a molecular readout, *M*, whose copy-number we denote by *n*_*M*_ . We assume that each graph, *G*, contains a subset of vertices, 𝒱_prod_(*G*) ⊂ 𝒱 (*G*), from which the system can produce *M* at a constant rate *r*. In addition, we assume that *M* undergoes first-order degradation, with rate *δn*_*M*_ . In this context, we can naturally define the steady-state level, SS(*x*), of *M* as the steady-state mean value of *n*_*M*_, which can be shown to equal to (Supplemental Information S1) [27, 28],

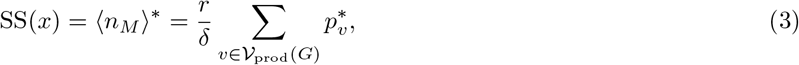

where, as discussed above, 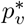 is the steady-state probability of the vertex *v*. For the chain models, we assume that 𝒱_prod_(*𝒞*_*N*_) = {*N*}; for the ladder models, we assume that 𝒱 _prod_(𝒟_*N*_) = {*U*_*N*_, *B*_*N*_}.

As discussed above, to obtain 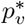 for each *v ∈* 𝒱 _prod_(*G*), Eqn. 1 tells us that the vector of steady-state probabilities, **p**^*∗*^, of the vertices in *G* lies in the kernel of ℒ (*G*). If *G* is *strongly connected* —that is, if every pair of vertices in *G* is connected by a path of directed edges—then one can show that dim ker ℒ (*G*) = 1, so that **p**^*∗*^ can be obtained by normalizing any vector, ***ρ*** *∈* ker *L*(*G*), by its coordinate sum [18]. The kinds of graphs we consider, 𝒞 _*N*_ and _*N*_, are strongly connected for all *N* . To calculate ***ρ*** in the numerical analyses we perform below, we obtained the singular value decomposition (SVD) of ℒ (*G*) and set ***ρ*** to the right singular vector corresponding to the zero singular value [29] (Materials and Methods).

On the other hand, we define the activation time of a system to produce one new molecule of *M* as a mean first-passage time (mFPT) on an augmented graph, *G*^+^, in which an additional “terminal” vertex is introduced to describe the production event. In particular, if *G* is a graph with vertices 𝒱 (*G*) = {1, …, *n*} and productive vertices 𝒱_prod_(*G*) ⊂ 𝒱 (*G*), then we define *G*^+^ as the graph on vertices 𝒱 (*G*^+^) = 𝒱 (*G*) ∪ {*M*} that is obtained by adding the edges *j* → *M*, for *j* ∈ 𝒱_prod_(*G*), each with label *ℓ*(*j* → *M*) = *r*. From here, we define the activation time as the mFPT,

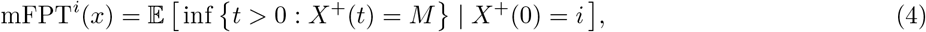

where *X*^+^(*·*) is the Markov process associated with *G*^+^, and *i* ∈ 𝒱 (*G*) is some initial vertex. This mFPT can be obtained from the matrix equation (Supplemental Information S1) [30],

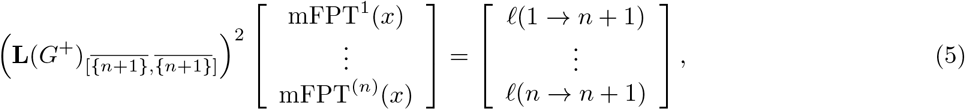

where we have introduced **L**(*G*^+^) = *−* ℒ (*G*^+^)^T^, and 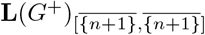 is the *n × n* sub-matrix of **L**(*G*^+^) obtained by removing the row and column corresponding to *M* ≡ *n* + 1. As such, for the purposes of the numerical analyses below, we solved this matrix equation by obtaining the QR decomposition of the above left-hand matrix [29] (Materials and Methods).

We also sought to obtain analytical formulas for SS(*x*) and mFPT^*i*^(*x*), both of which are accessible via the Matrix-Tree theorems [17–20]. To understand these formulas, we must first consider the *spanning forests* of a graph. For any graph Γ, a *spanning forest* of Γ is a subgraph, *F*, that (1) contains all the vertices in Γ (“spanning”); (2) contains no cycles, even when ignoring edge directions (“forest”); and (3) contains exactly one outgoing edge from each vertex other than a subset, ℛ (*F*) ⊂ 𝒱 (Γ), from which there are no outgoing edges. This subset of vertices are called the *roots* of *F* . If ℛ (*F*) consists of a single vertex, then *F* is a *spanning tree*. We denote the set of spanning forests of Γ rooted at *A* ⊂ 𝒱 (Γ) by Φ_*A*_(Γ).

Since there is exactly one outgoing edge from each non-root vertex in *F*, it is easy to see that *F* must contain a directed path of edges, *i → i*_1_ *→ · · · → i*_*k*_ *→ j*, from each vertex, *i ∉ R*(*F*), to *precisely* one root, *j ∈ R*(*F*). In particular, every root has a trivial path to itself, and if *F* is a spanning tree rooted at *R*(*F*) = {*j*}, then every vertex must have a path to *j* in *F* . Now, given *A* ⊂ 𝒱 (Γ) and *j* ∈ *A*, we denote by Φ_*A*:*i*⇝*j*_(Γ) ⊂ Φ_*A*_(Γ) the subset of spanning forests rooted at *A* in which there is a path from *i* to *j*. Here, *i* may either be any non-root vertex or *j*, so that *i* ∈ (𝒱(Γ) *\A*) ∪ {*j*}; in particular, we note that Φ_*A*:*j*⇝*j*_(Γ) = Φ_*A*_(Γ).

We are now ready to provide our formulas for SS(*x*) and mFPT^*i*^(*x*). For the former, the Matrix-Tree theorem tells us that the steady-state probability vector, **p**^*∗*^, is given by

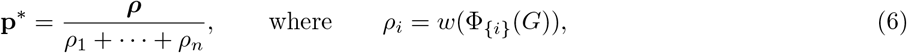

where *w*(*·*) is the *weight* function, which evaluates the sum of products of edge labels over each spanning forest in Φ_{*i*}_(*G*) [17–19]. More precisely, if ℋ is any collection of graphs, then

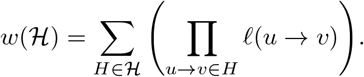

We can then use Eqn. 6 to calculate SS(*x*), per Eqn. 3. As for the mFPTs, we can use the All-Minors Matrix-Tree theorem to show that (Supplemental Information S1)

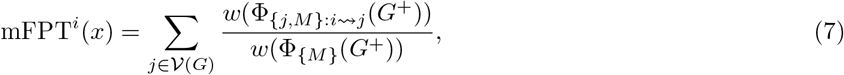

where, as described above, *G*^+^ is obtained by augmenting *G* with the terminal vertex, *M* [30].

Eqns. 6 and 7 demonstrate that analytical formulas for both quantities, in terms of the edge labels of the underlying graph, can be obtained through spanning tree/forest enumeration. However, such enumeration can be prohibitively expensive even for simple graphs. To circumvent this, we turn to a recurrence relation due to Chebotarev and Agaev [31], which can be written as follows. For any graph Γ with *m* vertices, let **Q**_*k*_(Γ) be the *m × m* matrix whose (*i, j*)-th entry is the weight of all spanning forests of Γ rooted at *m − k* vertices in which (1) *j* is a root and (2) there is a path of edges from *i* to *j*. In our notation, we can denote this as follows. If *i≠ j*, then the (*i, j*)-th entry of **Q**_*k*_(Γ) is given by,

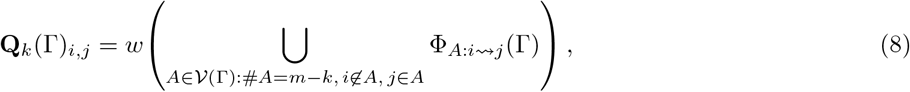

while the diagonal entries are given by

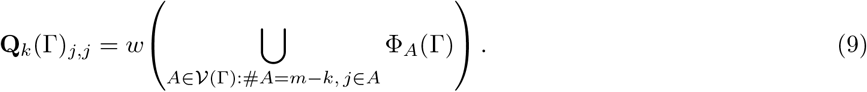

Then one can calculate **Q**_*k*_(Γ) for all *k* = 0, 1, … via the recurrence relation,

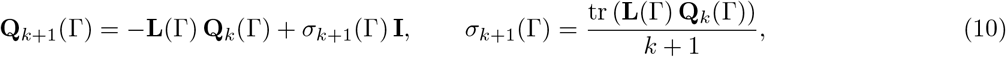

where **I** denotes the identity matrix and tr (*·*) denotes the trace, and the recurrence is initialized with **Q**_0_(Γ) = **I**.

Now, let us set Γ = *G*, in which case *m* = *n*. If we take *k* = *n −* 1 in Eqn. 9, we see that the only possible choice of *A* in the union is *A* = {*j*}. So we get (Eqn. 6)

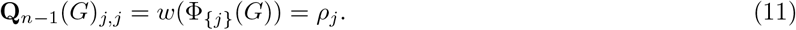

This means that, by calculating **Q**_*n−*1_(*G*) with Eqn. 10, we obtain *ρ*_*j*_, from which we can obtain SS(*x*) (Eqn. 3). For the mFPT (Eqn. 7), we require the doubly-rooted spanning forests of Γ = *G*^+^, whose weights contribute to **Q**_*n−*1_(*G*^+^). (Recall that *G*^+^ contains *m* = *n* + 1 vertices, due to the inclusion of the terminal vertex *M* .) In particular, we want to obtain the weight,

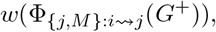

of all spanning forests rooted at {*j, M*} that contain a path from *i* to *j*, for all *j≠ M* . Setting Γ = *G*^+^ and *k* = *n −* 1 in Eqn. 8 yields

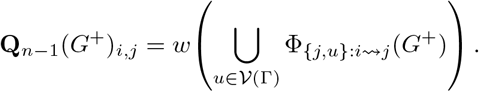

Now, suppose that *j* ≠ *M* . Then, since *M* is a terminal vertex and has no outgoing edges, every spanning forest of *G*^+^ must have *M* as a root. This means that, if *j* ≠ *M*, then the only choice of *u* for which the set in the above union is nonempty is *u* = *M* . Therefore, we get

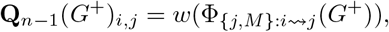

which provides the summands in the right-hand numerator of Eqn. 7. As for the denominator, setting *k* = *n* and *j* = *M* ≡ *n* + 1 in Eqn. 9 gives

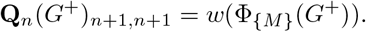

Putting these pieces together, we finally obtain

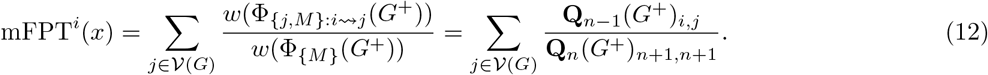

Therefore, evaluating **Q**_*n−*1_(*G*^+^) and **Q**_*n*_(*G*^+^) with Eqn. 10 gives us access to mFPT^*i*^(*x*). Taken together, Eqns. 10, 11 and 12 constitute a polynomial-time algorithm for exactly calculating SS(*x*) and mFPT^*i*^(*x*). Fig. S1 shows the agreement between the mFPT obtained using this recurrence and a set of simulated trajectories of *X*^+^(·) using the Gillespie algorithm [32].

To quantify decoupling between SS(*x*) and mFPT^*i*^(*x*), we consider normalised dynamic ranges for the two outputs, which we define as

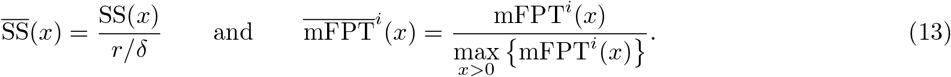

These definitions restrict 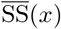 and 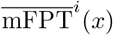 to between 0 and 1. Indeed, this restriction is self-evident for 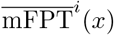; for 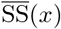, this stems from Eqn. 3, which tells us that

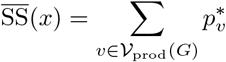

is the sum of steady-state probabilities for the subset of productive states, *v* ∈ 𝒱 _prod_(*G*). From here, we define the dynamic range of the two outputs as

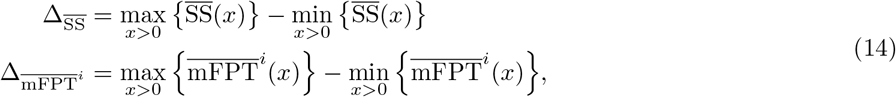

which also range between 0 and 1. We note that both 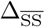 and 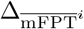 are relative dynamic ranges: the former is normalized by the theoretical maximum value of SS(*x*) (namely *r/δ*), while the latter is normalized by the actual maximum value of mFPT^*i*^(*x*). Therefore, both dynamic ranges account for changes in SS(*x*) and mFPT^*i*^(*x*) in proportion to these maximum values. We also note that perfect decoupling, in which only the steady-state level changes with input concentration, is obtained when 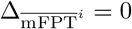 and 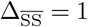.

## Results

### Decoupling under rate scale separation in *C*_2_ and *D*_2_

We start our analysis by considering the two-state chain model, 𝒞_2_, which is equivalent to the “random telegraph” model of transcriptional bursting (Fig. 1E) [14, 33, 34]. Here, vertex 1 represents an inactive state, and vertex 2 represents an active state that can produce the molecular readout, *M*, so that 𝒱_prod_(𝒞_2_) = {2} . The steady-state level is given by (Eqns. 3 and 6; Supporting Information S1) [21, 27]:

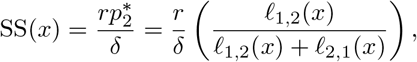

whereas the activation time is given by (Eqn. 7)

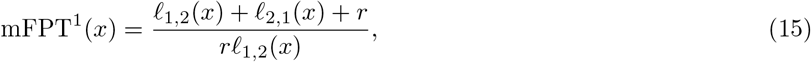

where we have set the initial vertex to *i* = 1. Notice that, upon sending *r →* ∞, we have

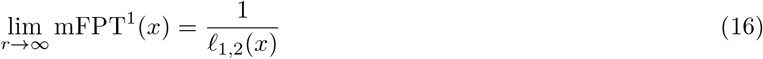

From this, we can see that if the ligand acts only on the edge 2 → 1, such that *ℓ*_1,2_(*x*) = *ℓ*_1,2_ does not depend on *x*, and the production rate is large (*r* → ∞), then mFPT^1^(*x*) remains constant while the steady-state level, SS(*x*), changes with *x*. Thus, in this regime, the ligand may only affect the steady-state level, but not the activation time, leading to decoupling. This simple model immediately suggests that decoupling can be easily achieved if the ligand only acts on the deactivation transition, as long as production from vertex 2 is sufficiently fast such that, upon reaching vertex 2, the system produces *M* before transitioning back to vertex 1. We call this *rate scale separation*. If the ligand acts on the edge 1 *→* 2 as well, so that *ℓ*_1,2_(*x*) also depends on *x*, whether decoupling is possible depends on the functional forms of *ℓ*_1,2_(*x*) and *ℓ*_2,1_(*x*).

In order to examine the implications of assuming a specific mechanism by which the bound ligand affects the system, we next considered the corresponding ladder model, *D*_2_ (Fig. 1F). Here, recalling that 𝒱_prod_(*D*_2_) = {*U*_2_, *B*_2_}, the steady-state level of *M* is defined as (Eqns. 3 and 6)

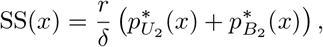

and we can similarly use Eqn. 7 to derive the activation time, 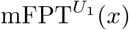, with initial vertex *i* = *U*_1_.

The ligand can either promote or hinder production of the readout; in this model, this is determined by the values of the regulatory factors, *γ*_1,2_ and *γ*_2,1_. The former measures the strength with which the ligand regulates the forward transition, *B*_1_ *→ B*_2_; the latter measures the strength with which the ligand regulates the backward transition, *B*_2_ → *B*_1_. To simplify our analysis, we will assume that the ligand acts on only one of the two transitions, and that it *promotes* readout production, thus acting as an *activator*. Mathematically, this means that either *γ*_1,2_ *>* 1 and *γ*_2,1_ = 1, or *γ*_1,2_ = 1 and *γ*_2,1_ *<* 1. Using the mathematical machinery laid out in the previous section, it can be shown that, in this regime, SS(*x*) increases monotonically with *x* and 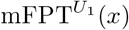 decreases monotonically with *x* (Supporting Information S2). In this case, analytical expressions for the corresponding dynamic ranges can be obtained by comparing the values of 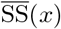 and 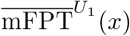 at *x* = 0 and *x →* ∞. In particular, when the forward transition is regulated by the ligand (*γ*_1,2_ *>* 1, *γ*_2,1_ = 1), we obtain the following dynamic ranges:

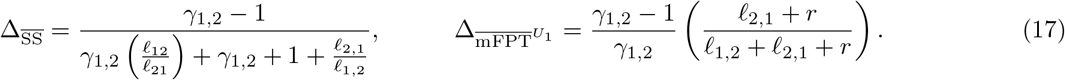

Meanwhile, when the ligand regulates the backward transition (*γ*_1,2_ = 1, *γ*_2,1_ *<* 1), we obtain,

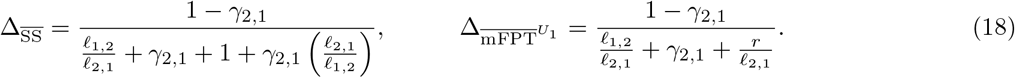

The expressions in Eqn. 18 reveal that, when the backward transition is regulated by the ligand, 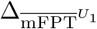 tends to zero as *r →* ∞, while 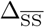 does not depend on *r*. However, if the forward transition is regulated by the ligand (Eqn. 17), then taking the same limit causes 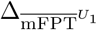 to converge to a finite nonzero value, namely (*γ*_1,2_ *−* 1)*/γ*_1,2_. Therefore, a similar result to what we found for 𝒞_2_ holds for 𝒟_2_: the two outputs can be decoupled if the ligand regulates the backward transition, *B*_2_ *→ B*_1_, and the production rate, *r*, is large. Notice that, in both equations, 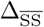 and 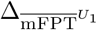 may both tend to zero in other limiting regimes, e.g., *ℓ*_1,2_*/ℓ*_2,1_ *→* ∞; this does not correspond to our definition of decoupling, but rather a trivial case of unresponsiveness of the system.

### Decoupling under rate scale separation in *D*_3_

Molecular systems in various biological settings often transition through multiple states before producing the molecular readout [9, 14, 15, 35, 36]. We thus asked what happens in the ladder model *D*_3_.

To begin with a simplified setting, we assumed *ℓ*_1,2_ = *ℓ*_2,1_ and *ℓ*_2,3_ = *ℓ*_3,2_, and that the ligand regulates exactly one of the four transitions, *B*_1_ → *B*_2_, *B*_2_*→ B*_1_, *B*_2_→ *B*_3_, and *B*_3_ → *B*_2_, promoting readout production. In this case, there are four possibilities, which are enumerated in Table 1. For each of these regimes, it can be shown that SS(*x*) and 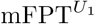 (*x*) are both monotonic in *x* (Supporting Information S2) and analytical expressions for the dynamic ranges of the corresponding normalised quantities can be obtained, analogously to the 𝒟 _2_ model.

**Table 1.** Dynamic ranges of 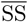 and 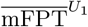 in 𝒟_3_, for regulatory regimes that promote readout production through the regulation of one transition.

| regulatory regimes | | $\Delta_{\overline{SS}}$ | $\Delta_{\overline{mFPT}^{U_1}}$ |
| --- | --- | --- | --- |
| 1.I | $\gamma_{1,2} > 1$ | $\frac{\gamma_{1,2} - 1}{3(2\gamma_{1,2} + 1)}$ | $\frac{\gamma_{1,2} - 1}{\gamma_{1,2}} \left( \frac{\beta + \alpha + \alpha\beta}{\alpha\beta + 2\beta + 3\alpha} \right)$ |
| 1.II | $\gamma_{2,1} < 1$ | $\frac{1 - \gamma_{2,1}}{3(\gamma_{2,1} + 2)}$ | $(1 - \gamma_{2,1}) \left( \frac{\alpha + \beta}{\alpha\beta + 2\beta + 3\alpha} \right)$ |
| 1.III | $\gamma_{2,3} > 1$ | $\frac{2(\gamma_{2,3} - 1)}{3(\gamma_{2,3} + 2)}$ | $\frac{\gamma_{2,3} - 1}{\gamma_{2,3}} \left( \frac{2(\alpha + \beta)}{\alpha\beta + 2\beta + 3\alpha} \right)$ |
| 1.IV | $\gamma_{3,2} < 1$ | $\frac{2(1 - \gamma_{3,2})}{3(2\gamma_{3,2} + 1)}$ | $(1 - \gamma_{3,2}) \left( \frac{2\alpha}{\alpha\beta + 2\beta + 3\alpha} \right)$ |

The normalised dynamic ranges for the four parametric regimes are summarized in Table 1 below, where we have introduced the dimensionless parameters,

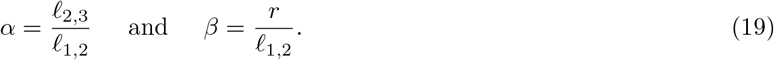

We first notice that, in almost all the cases, having a large production rate (*r →* ∞) is no longer sufficient to get decoupling. The only case in which this is sufficient is case 1.IV, where only the last backward transition, *B*_3_ *→ B*_2_, is regulated. Here, we may rewrite 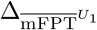 as,

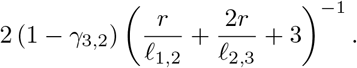

This is in line with the findings for 𝒟_2_ in the previous section, and can be understood intuitively as follows. When *r* is sufficiently large, every time the system reaches *U*_3_ or *B*_3_, it will rapidly proceed to *M* without backtracking to vertex *U*_2_ or *B*_2_, respectively. Therefore, the mFPT from *U*_1_ to *M* can be approximated as the mFPT from *U*_1_ to *U*_3_ or *B*_3_, whichever is reached first. Now, if the ligand does not regulate any transition other than *B*_3_ *→ B*_2_, the dynamics with which the system proceeds to *U*_3_ or *B*_3_ in the limits of zero or infinite ligand concentration, respectively, are the same on average. Since the mFPT is monotonic in *x* (Supplemental Information S2), this implies that the mFPT does not change with *x*. Therefore, in case 1.IV, a large production rate is sufficient for decoupling.

In addition, we notice that for cases 1.II, 1.III and 1.IV, but not 1.I, there exists a different parametric regime in which 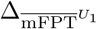 tends to zero but 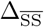 does not, thus giving rise to decoupling: *α, β* ≫ 1, which we may rewrite in terms of rates as *ℓ*_2,3_, *r* ≫ *ℓ*_1,2_. In particular, we note that, in cases 1.II and 1.III, it is not sufficient to have *either α* ≫ 1 *or β* ≫ 1; rather, both *α* and *β* must be large. In this rate-scale-separated regime, the first forward transition in the absence of ligand, *U*_1_ *→ U*_2_, is much slower than the second, *U*_2_ *→ U*_3_, as well as the production transitions, *U*_3_ *→ M* and *B*_3_ *→ M* . In addition, when *γ*_1,2_ = 1 (as in cases 1.II, 1.III, and 1.IV), the first forward transition in the presence of ligand, *B*_1_ *→ B*_2_, is also much slower than *U*_2_ *→ U*_3_, *B*_2_ *→ B*_3_, *U*_3_ *→ M*, and *B*_3_ *→ M* . We hypothesised that, in this case, a partitioning of the graph arises where the slow rate, namely *ℓ*_1,2_, completely determines the activation time, while the fast rates dictate the dynamic range of the steady-state level. To test this hypothesis, we examined 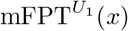 for cases 1.II, 1.III and 1.IV, when *x* = 0. For all three cases, the activation time is given by (Eqn. 12 and setting *x* = 0),

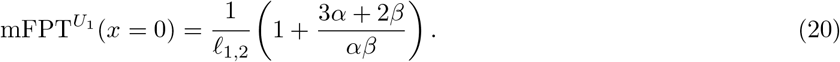

Here, imposing *α, β* ≫ 1 yields an activation time that depends merely on *ℓ*_1,2_. Since, as shown above, 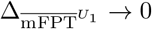 in this parametric regime for cases 1.II, 1.III and 1.IV, this means that 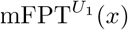 depends merely on *ℓ*_1,2_ for all *x*. Therefore, under rate scale separation, the activation time depends entirely on the *slower, unregulated* forward transition.

Meanwhile, the expression for the steady-state level, SS(*x*), depends on the case, as well as which parametric limits are applied to realise the condition that *α* and *β* should be large. For instance, in the simple setting in which we send *ℓ*_1,2_ *→* 0, the steady-state level in case 1.III is given by (Eqn. 6 and symbolic limit for *ℓ*_1,2_ *→* 0),

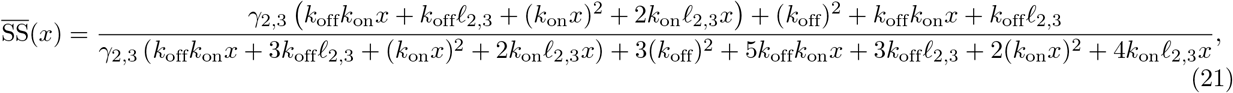

which depends on the fast rate *ℓ*_2,3_, and the regulatory factor, *γ*_2,3_. In contrast, in case 1.II, in which the slow backward transition *B*_2_ *→ B*_1_ is regulated, we have (Eqn. 6 and symbolic limit for *ℓ*_1,2_ *→* 0),

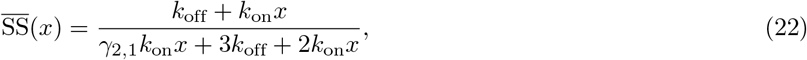

which depends on the regulatory factor, *γ*_2,1_, but not the fast rate, *ℓ*_2,3_.

Next, we aimed to assess whether rate scale separation can still give rise to decoupling if we relax some of the assumptions made in the above analysis, namely that *ℓ*_1,2_ = *ℓ*_2,1_ and *ℓ*_2,3_ = *ℓ*_3,2_. We found that, upon relaxing these assumptions, the expressions for 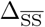 and 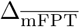 become substantially more complicated; therefore, we resorted to numerical optimization to search for parameter regimes that lead to decoupling, focusing on case 1.III (*γ*_2,3_ *>* 1; Fig. 2A). We defined a *coupling score*,

**Fig. 2.**
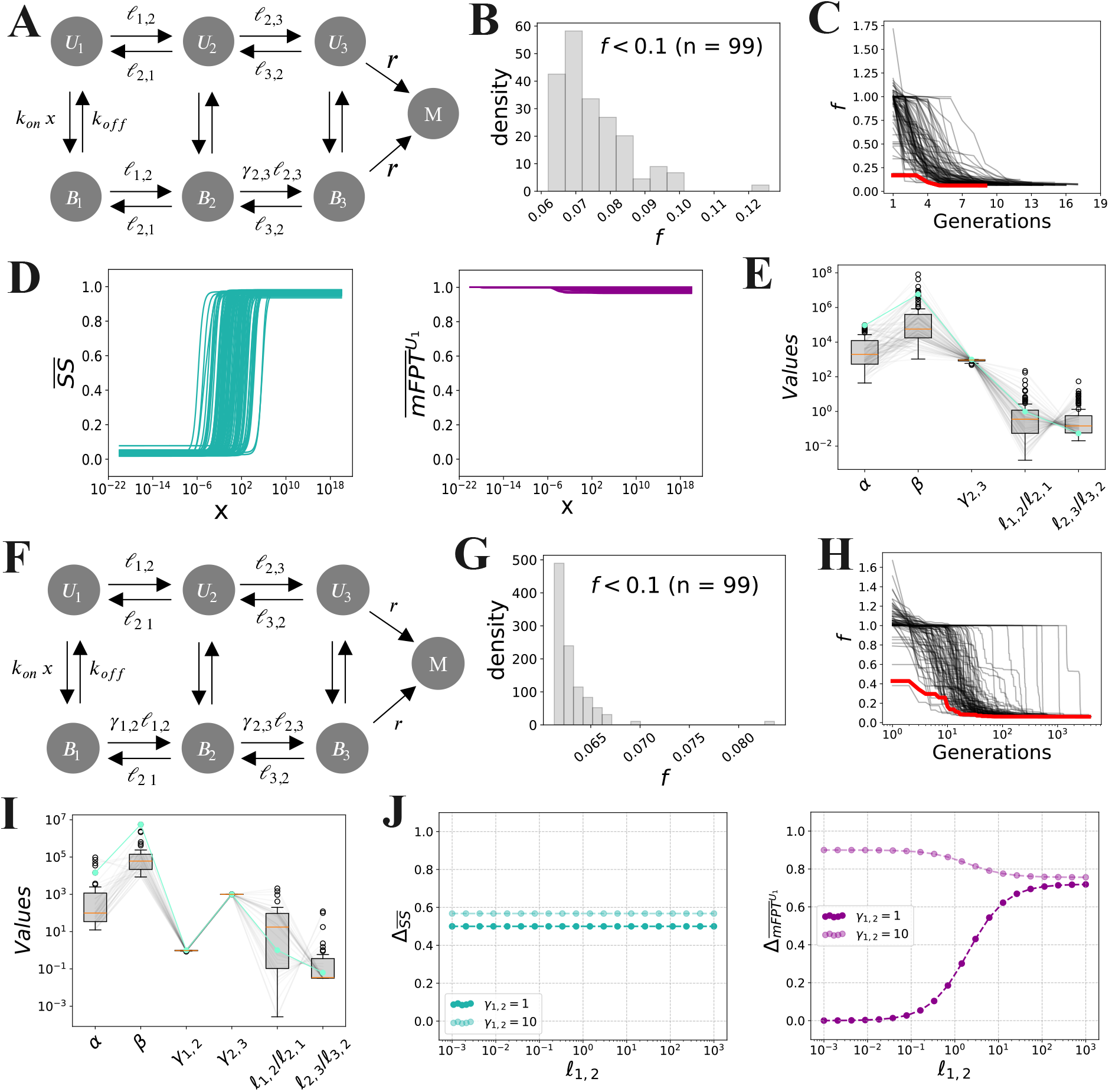
Decoupling in the ladder model, *D*_3_, for cases 1.III (A–E) and a generalization of case 2.I (F–J). **(A)** Schematic of *D*_3_ with regulation of *B*_2_ *→ B*_3_ (case 1.III). **(B)** Distribution of coupling scores after termination of the PSO. Each PSO run was terminated whenever *f <* 0.05 for more than 5 consecutive generations. **(C)** Evolution of *f* over each PSO run. The red curve represents the “best” optimized parameter set with the smallest value of *f* . **(D)** Input-output responses of optimized parameter sets for which *f <* 0.05 and the steady-state level increases monotonically with *x*. **(E)** Distributions of parameter values corresponding to panel D, with *α* = *ℓ*_2,3_*/ℓ*_1,2_ and *β* = *r/ℓ*_1,2_. The green curve represents the best parameter set. **(F)** Schematic of 𝒟 _3_ with regulation of *B*_1_ *→ B*_2_ and *B*_2_ *→ B*_3_ (generalization of case 2.I). **(G)** Distribution of coupling scores after termination of the PSO. Each PSO run was terminated after 23 hours of computation time. **(H)** Evolution of *f* over each PSO run. The red curve represents the best parameter set. **(I)** Distributions of optimized parameter values for which *f <* 1 and the steady-state level increases monotonically with *x*. **(J)** Normalized dynamic ranges for two families of parameter sets, with the parameters set as follows: *ℓ*_2,3_ = *ℓ*_3,2_ = 10*δ, k*_off_ = *δ, k*_on_ = *δ/* (1 c.u.), *r* = 10*δ, γ*_2,3_ = 10, *γ*_1,2_ = 1 or 10, and *ℓ*_1,2_ = *ℓ*_2,1_ varied over a logarithmic range. The dots represent numerical computations (Materials and Methods), and the dashed lines represent the formulas in Table 2 (case 2.I).

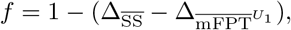

which describes the extent of coupling between the steady-state level and activation time. Perfect decoupling corresponds to *f* = 0, corresponding to a maximum steady-state dynamic range 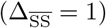 and a minimum activation time dynamic range 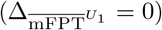. We then searched for parameter sets that yield a small value of *f* by using a Particle Swarm Optimization (PSO) algorithm [37, 38] (Materials and Methods). Briefly, this algorithm begins with a collection (or “swarm”) of parameter sets (or “particles”), and iteratively updates each particle’s position and velocity according to the values of the objective function (here, *f*) across the swarm. Over successive generations, the swarm collectively converges to the optimal solution(s) with respect to the objective function.

We allowed the rates, *ℓ*_*i,j*_, *k*_off_, and *r*, to lie within a large parameter range, namely [10^*−*4^, 10^4^] in units of *δ*. The binding rate constant, *k*_on_, was also assumed to lie in the range [10^*−*4^, 10^4^], but in units of *δ/* (1 c.u.), where “c.u.” denotes the concentration units used for *x*. Finally, the dimensionless regulatory parameter, *γ*_2,3_, was constrained to lie in the range [1, 10^3^]. Each parameter was restricted to lie within these ranges throughout the optimization. We ran 100 independent optimization runs, each starting from a different random initial condition. Figs. 2B and C show that more than 90% of the runs converged to a final coupling score of *f <* 0.1, strongly suggesting that this procedure effectively minimizes the objective function.

Among these successful optimization runs, we filtered out the optimised parameter sets for which the steady-state level is constant in *x*, and visualised the normalised steady-state level and activation time for the remaining parameter sets (Fig. 2D). This revealed that, while the steady-state level increases monotonically with *x* (Fig. 2D, left), the mFPT does not change significantly within the same concentration range (Fig. 2D, right), demonstrating near-perfect decoupling. In line with our analytical results, we found that (1) the values of *α* and *β* are both large, and (2) the second forward transition, *B*_2_ → *B*_3_, is strongly promoted by the ligand, with *γ*_2,3_ almost always reaching its maximum value of 10^3^ (Fig. 2E). This strongly suggests that our optimization procedure is identifying parameter regimes that achieve decoupling via rate scale separation.

Notably, we also found that *ℓ*_3,2_ often exceeds *ℓ*_2,3_, albeit to an extent far less than *r*; for instance, the best parameter set over all optimisation runs (Fig. 2E, green) exhibited *ℓ*_3,2_*/ℓ*_2,3_ ∼ 10 and *β/α* = *r/ℓ*_2,3_ ∼ 10^5^. We also found a broad distribution of values for *ℓ*_1,2_*/ℓ*_2,1_. This indicates that the assumptions that *ℓ*_1,2_ = *ℓ*_2,1_ and *ℓ*_2,3_ = *ℓ*_3,2_ are not necessary for decoupling via rate scale separation. Indeed, combining these results with the analytical formulas in Table 1 reveals that *ℓ*_3,2_ *> ℓ*_2,3_ *strengthens* decoupling, by increasing the steady-state dynamic range, 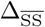 . Specifically, a regulatory regime in case 1.III that satisfies *ℓ*_1,2_ = *ℓ*_2,1_ and *ℓ*_2,3_ = *ℓ*_3,2_ has a steady-state dynamic range of,

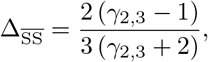

which, in the limit of large *γ*_2,3_, converges to a maximum value of 2*/*3. This is because the steady-state level of the readout is nonzero, even when ligand is absent. In particular, if *ℓ*_1,2_ = *ℓ*_2,1_ and *ℓ*_2,3_ = *ℓ*_3,2_, then it is easy to apply Eqn. 6 to find that the steady-state probabilities of the six states at zero ligand concentration are given by,

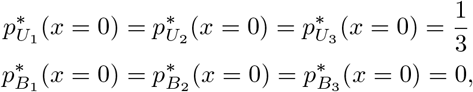

whereas, in the limit of large ligand concentration, we instead have,

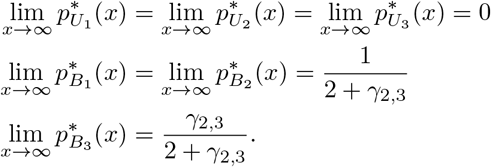

Therefore, if *γ*_2,3_ ≫ 1, then we have,

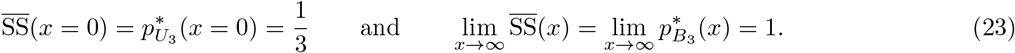

Since 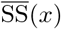 is monotonic in *x* in this case, this means that 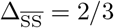. On the other hand, allowing for *ℓ*_3,2_ *> ℓ*_2,3_ decreases the steady-state probability of *U*_3_ relative to those of *U*_1_ and *U*_2_ at *x* = 0, and therefore decreases the steady-state level of the readout. This illustrates how asymmetry in the transition rates acts in concert with the regulatory factor, *γ*_2,3_, to modulate the steady-state dynamic range, whereas *α* and *β* together dictate the mFPT dynamic range, thus inducing decoupling.

We next proceeded to extend our analysis to regulatory regimes in which the ligand regulates *two* transitions. There are 16 such possible regulatory regimes, depending on the choice of ligand-regulated transitions and whether each corresponding regulatory factor, *γ*_*i,j*_, is either greater than or less than 1. We can categorize these regimes into two classes: eight *coherent* regimes, in which the ligand consistently promotes or hinders transitioning towards the productive state (e.g., *γ*_1,2_ *>* 1 and *γ*_2,3_ *>* 1); and eight *incoherent* regimes, in which the ligand simultaneously promotes and hinders transitioning towards the productive state (e.g., *γ*_1,2_ *>* 1 and *γ*_2,3_ *<* 1) [23]. We first restricted our attention to the four *coherent* regulatory regimes in which the ligand consistently *promotes* transitioning towards the productive state; these regimes are enumerated in Table 2. For each of these regimes, it can be shown that SS(*x*) and 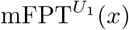 increase and decrease monotonically with *x*, respectively (Supporting Information S2), and we can obtain analytical expressions for the normalised dynamic ranges of these quantities, as before. These expressions are given in Table 2.

**Table 2.** Dynamic ranges of 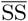 and 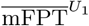 in 𝒟, for regulatory regimes that promote readout production through the regulation of two transitions.

| regulatory regimes | | $\Delta_{\overline{\text{SS}}}$ | $\Delta_{\overline{\text{mFPT}}^{U_1}}$ |
| --- | --- | --- | --- |
| 2.I | $\gamma_{1,2} > 1, \gamma_{2,3} > 1$ | $\frac{2\gamma_{1,2}\gamma_{2,3} - \gamma_{1,2} - 1}{3(\gamma_{1,2}\gamma_{2,3} + \gamma_{1,2} + 1)}$ | $\frac{\alpha\beta\gamma_{2,3}(\gamma_{1,2} - 1) + (\alpha + \beta)(2\gamma_{1,2}\gamma_{2,3} - \gamma_{1,2} - 1)}{\gamma_{1,2}\gamma_{2,3}(\alpha\beta + 3\alpha + 2\beta)}$ |
| 2.II | $\gamma_{1,2} > 1, \gamma_{3,2} < 1$ | $\frac{2\gamma_{1,2} - \gamma_{1,2}\gamma_{3,2} - \gamma_{3,2}}{3(\gamma_{1,2}\gamma_{3,2} + \gamma_{1,2} + \gamma_{3,2})}$ | $\frac{(\alpha\beta + \beta)(\gamma_{1,2} - 1) + \alpha(2\gamma_{1,2} - \gamma_{1,2}\gamma_{3,2} - \gamma_{3,2})}{\gamma_{1,2}(\alpha\beta + 3\alpha + 2\beta)}$ |
| 2.III | $\gamma_{2,1} < 1, \gamma_{2,3} > 1$ | $\frac{2\gamma_{2,3} - \gamma_{2,1} - 1}{3(\gamma_{2,1} + \gamma_{2,3} + 1)}$ | $\frac{(\alpha + \beta)(2\gamma_{2,3} - \gamma_{2,1} - 1)}{\gamma_{2,3}(\alpha\beta + 3\alpha + 2\beta)}$ |
| 2.IV | $\gamma_{2,1} < 1, \gamma_{3,2} < 1$ | $\frac{2 - \gamma_{2,1}\gamma_{3,2} - \gamma_{3,2}}{3(\gamma_{2,1}\gamma_{3,2} + \gamma_{3,2} + 1)}$ | $\frac{\beta(1 - \gamma_{2,1}) + \alpha(2 - \gamma_{2,1}\gamma_{3,2} - \gamma_{3,2})}{\alpha\beta + 3\alpha + 2\beta}$ |

Although these expressions are more complicated, we still see that imposing rate scale separation, by setting *α, β* ≫ 1, causes 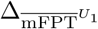, but not 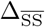, to tend to zero, as long as the slow, forward transition (*B*_1_ *→ B*_2_) is not regulated (cases 2.III and 2.IV). Moreover, for each of these two cases, it is easy to directly compare the expressions for 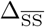 and 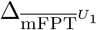 to the expressions that arise when only one transition is regulated (cases 1.II and 1.III for 2.III, and cases 1.II and 1.IV for 2.IV), to see that 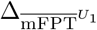 increases and 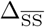 decreases when a second transition is regulated. As such, introducing a second regulated transition enhances decoupling.

We next assessed the relevance of the constraints *ℓ*_1,2_ = *ℓ*_2,1_ and *ℓ*_2,3_ = *ℓ*_3,2_, again using the numerical optimization procedure outlined above. We aimed to minimize the coupling score, *f*, for a generalized version of case 2.I (Fig. 2F), in which *γ*_1,2_ and *γ*_2,3_ can both assume any value within the range, [10^*−*3^, 10^3^] . Similarly to the optimization for case 1.III, we found that more than 90% of the optimization runs converge to a coupling score of *f <* 0.1 (Fig. 2G), albeit with a larger number of generations (Fig. 2H), as expected by the increased dimensionality of the parameter space. Notably, we found that the optimal parameter sets exhibit significant rate scale separation, *α* ≫ 1 and *β* ≫ 1, as well as values of *γ*_1,2_ ≈ 1, representing little to no regulation of the slow forward transition *B*_1_ → *B*_2_, and *γ*_2,3_ close to the maximum value of 10^3^ (Fig. 2I). This strongly suggests that, to attain decoupling, the algorithm is effectively reducing this generalization of case 2.I to a regulatory regime in case 1.III, in which *B*_1_ *→ B*_2_ is unregulated. In addition, we found that the ratios *ℓ*_1,2_*/ℓ*_2,1_ and *ℓ*_2,3_*/ℓ*_3,2_ follow similar distributions as in case 1.III (Fig. 2E), with *ℓ*_3,2_*/ℓ*_2,3_ ∼ 10 in the best parameter set (Fig. 2I, green). This, again, reflects the fact that increasing *ℓ*_3,2_*/ℓ*_2,3_ decreases 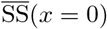, and therefore increases 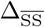, thus strengthening decoupling.

The equations in Table 1 and Table 2 and the numerical results in Fig. 2 show that a large rate scale separation can give rise to decoupling. To assess whether decoupling can be achieved in a more constrained scenario, we examined a family of example parameter sets in case 1.III with *γ*_2,3_ = 10, *ℓ*_2,3_ = *ℓ*_3,2_ = *r* = 10*δ, k*_off_ = *δ*, and *k*_on_ = *δ/* (1 c.u.), and plotted 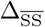 and 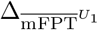 while varying *ℓ*_1,2_ = *ℓ*_2,1_ over several orders of magnitude (Fig. 2J, dark colours). Note that, in this case, *α* = *β*. Here, we found that, while 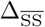 did not significantly vary with *ℓ*_1,2_, we could decrease 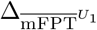 to as small as 0.1 by setting *α* = *β* ≈ 10, suggesting that values of *α, β*, and *γ*_2,3_ much less than those reported in Fig. 2E can also give rise to significant decoupling. Meanwhile, we found that the same family of parameter sets but with *γ*_1,2_ = 10, which instead fall under case 2.I, did not exhibit significant decoupling for any choice of *ℓ*_1,2_ = *ℓ*_2,1_ (Fig. 2J, light colours), consistent with our observations in Fig. 2I.

In summary, our numerical and analytical results demonstrate that rate scale separation, when paired with a lack of regulation of the slower forward transition, *B*_1_ *→ B*_2_, gives rise to decoupling in the ladder model, 𝒟_3_. In order to check whether this same mechanism enables decoupling in larger models, we performed a similar analysis of 𝒟_6_ and found that, there too, decoupling can arise when (1) the system exhibits a block of transitions that are slower than a subsequent block of transitions, and (2) regulation occurs along one or more of the faster forward transitions (Fig. S9A–C). Such separations of timescales have been widely recognised in the setting of gene regulation, where changes in chromatin state are typically measured to proceed on slower timescales than TF binding and the reactions that comprise the polymerase cycle [33, 39–41]. This suggests that decoupling in gene regulation could arise when activating TFs do not accelerate the slow chromatin opening transitions, but rather work on other subsequent steps.

### Decoupling due to incoherent regulation, in the absence of rate scale separation

We then asked whether there are alternative regulatory mechanisms that can yield decoupling when all transitions operate on similar timescales. Recently, we and others have argued that TFs may act on multiple steps of a gene-regulatory mechanism in an incoherent fashion, simultaneously promoting and hindering transcription [21, 23]. In the light of this, we hypothesized that such incoherent regulation may be an alternative way to maintain a constant activation time, perhaps by counterbalancing the effects of promoting progression towards the productive state through certain transitions by hindering this progression along other transitions, all while allowing for a change in the steady state.

To examine how decoupling might arise when the transition rates are constrained to be similar, we first considered the extreme scenario in which *ℓ*_1,2_ = *ℓ*_2,1_ = *ℓ*_2,3_ = *ℓ*_3,2_. In this case, assuming that the ligand acts only on the forward transitions *B*_1_ *→ B*_2_ and *B*_2_ *→ B*_3_ (i.e., *γ*_1,2_, *γ*_2,3_ ≠ 1 and *γ*_2,1_ = *γ*_3,2_ = 1), it can be shown that decoupling cannot be achieved in any coherent regulatory regime in which *γ*_1,2_ ≥ 1 and *γ*_2,3_ ≥ 1. In particular, it can be shown that, in this regime (Supporting Information S3),

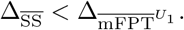

This implies that, whenever *γ*_1,2_ ≥ 1 and *γ*_2,3_ ≥ 1, the coupling score, *f*, must be greater than one. Therefore, if the ligand is assumed to promote one of the forward transitions *and ℓ*_1,2_ = *ℓ*_2,1_ = *ℓ*_2,3_ = *ℓ*_3,2_, decoupling may only be achieved if the ligand *hinders* the other forward transition.

Now, if we allow for incoherent regulation by the ligand (e.g., *γ*_1,2_ *>* 1 and *γ*_2,3_ *<* 1), then 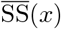 and 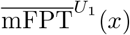 are not necessarily monotonic in *x* (Supplemental Information S2). However, we can still characterise conditions under which 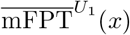 is not necessarily constant in *x*, but satisfies the weaker condition,

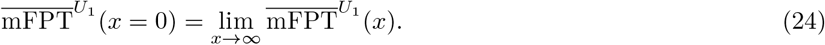

In particular, we can show that, if we set *ℓ*_1,2_ = *ℓ*_2,3_ = *ℓ*_2,1_ = *ℓ*_3,2_ = *r* and allow for regulation along *B*_1_ *→ B*_2_ and *B*_2_ *→ B*_3_ (*γ*_1,2_, *γ*_2,3_ ≠ 1 and *γ*_2,1_ = *γ*_3,2_ = 1), then we have,

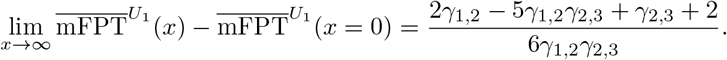

Equating this to zero and solving for either *γ*_1,2_ or *γ*_2,3_, we obtain,

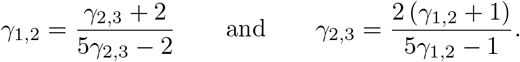

From here, it is easy to see that *γ*_1,2_ *>* 1 if, and only if, *γ*_2,3_ *<* 1. This implies that any coherent regulatory regime in which *γ*_1,2_, *γ*_2,3_ *>* 1 or *γ*_1,2_, *γ*_2,3_ *<* 1 cannot satisfy Eqn. 24, and therefore cannot exhibit a constant 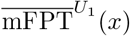 in *x*. This demonstrates that, in the extreme scenario where *ℓ*_1,2_ = *ℓ*_2,1_ = *ℓ*_2,3_ = *ℓ*_3,2_ = *r*, incoherent regulation is necessary to achieve 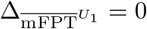.

To pursue a more comprehensive analysis of decoupling in the situation where the transitions proceed with similar rates, we again turned to numerical optimisation. In particular, we adapted our PSO approach to incorporate a “Rate Scale Constraint” (RSC) that constrains the ratio between each pair of transition rates, as follows:

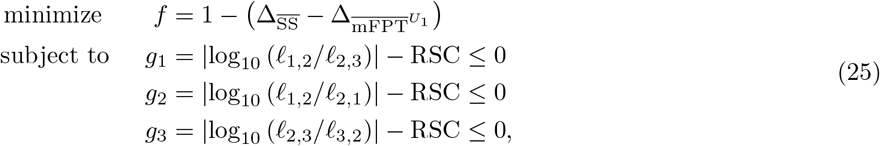

where RSC is some positive constant. The smaller RSC is, the more similar *ℓ*_1,2_, *ℓ*_2,1_, *ℓ*_2,3_, and *ℓ*_3,2_ tend to be. Within the PyMoo optimisation framework that we used to perform PSO [38], inequality constraints are handled as penalties to the objective function; as such, we independently confirmed that all solutions obtained from the PSO do satisfy the constraints given in Eqn. 25 (Fig. S2C, left). We emphasise that we did not impose any constraints on the ligand’s regulatory mode, in principle allowing for both coherent and incoherent regulation.

We first focused on the case where ligand may regulate the two forward transitions, *B*_1_ *→ B*_2_ and *B*_2_ *→ B*_3_ (so that *γ*_1,2_ and/or *γ*_2,3_ may be distinct from 1, and *γ*_2,1_ = *γ*_3,2_ = 1). We ran PSO with six different values for RSC: 0.005, 0.05, 0.5, 1, 2, and 3. As before, to increase the probability of finding global optima, we performed 100 replicates of the optimization, each starting from a different random initial condition. The convergence of each replicate and fulfillment of the RSC constraints are shown in Figs. S2A and S2C (left), respectively. Consistent with the analyses throughout the manuscript, we considered only activating responses, where the steady-state response increases with TF concentration.

Fig. 3A shows that, as RSC → 0, we obtained a larger minimum coupling score, suggesting that it is more difficult to obtain decoupling under rate scale constraint. When examining the corresponding input-output curves, we observed some dependence of the normalised mFPT on the ligand concentration (Fig. 3B, bottom). Yet, we observed that the increase in the coupling score was mostly determined by a smaller dynamic range in the steady-state level, arising from a nonzero basal steady-state level at zero input concentration (Fig. 3B, top). This is consistent with our previous reasoning used to derive Eqn. 23: the closer *ℓ*_1,2_*/ℓ*_2,1_ and *ℓ*_2,3_*/ℓ*_3,2_ are to 1, the closer the normalised steady-state level at zero ligand concentration is to 1*/*3, which is indeed the value of 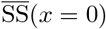 that we observe in our optimisation results (Fig. 3B, top).

**Fig. 3.**
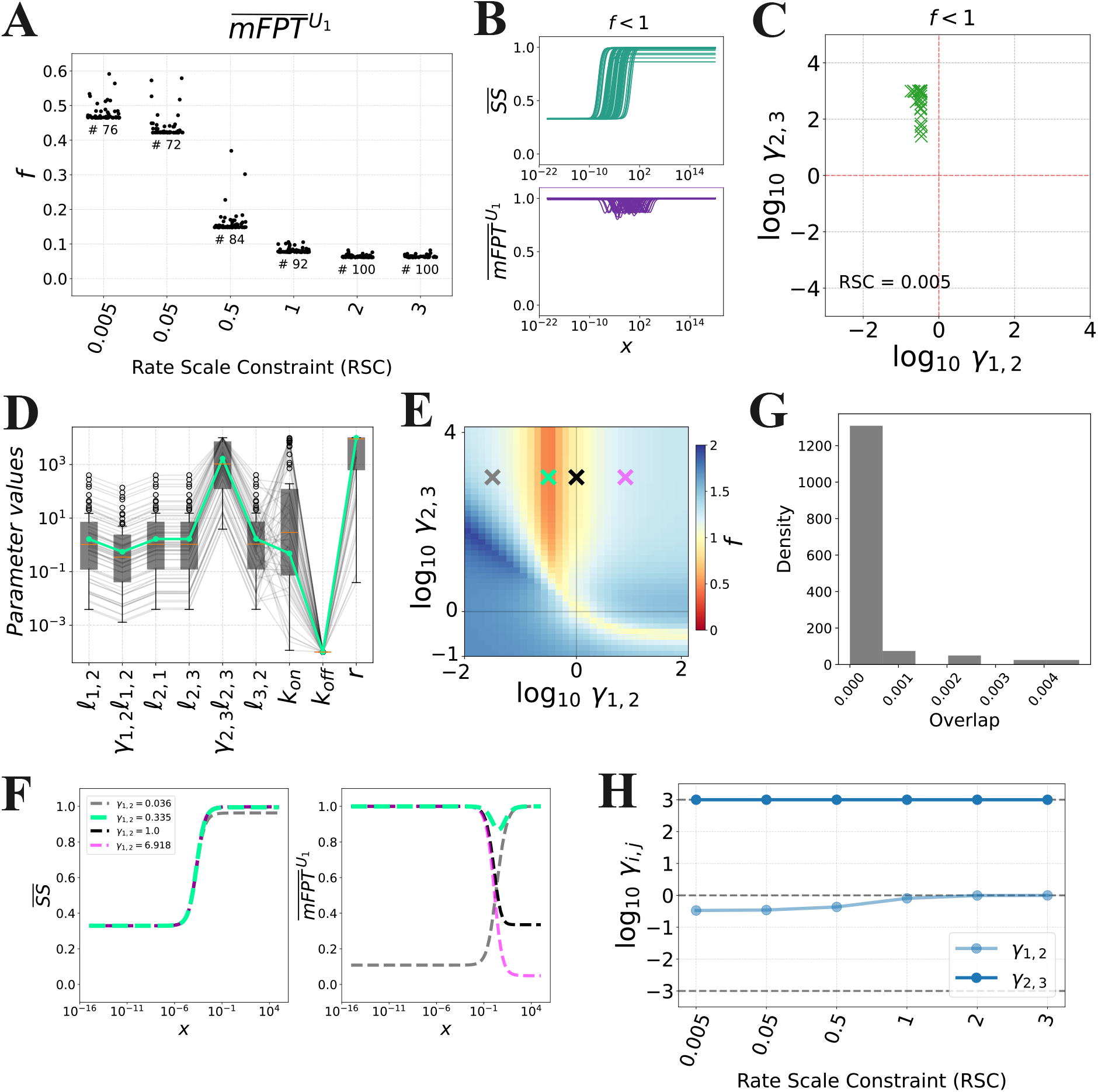
Decoupling under rate scale constraint arises under incoherent regulation. **(A)** Distributions of the coupling score, *f*, obtained from optimization with various rate scale constraints (RSC). **(B)** Input-output responses corresponding to the parameter sets obtained from optimization with RSC = 0.005. **(C)** Values of *γ*_1,2_ and *γ*_2,3_ for optimal parameter sets for RSC = 0.005. **(D)** Values of the parameter sets from each optimization run for RSC = 0.005. The green line represents the best parameter set. Each parameter is plotted in units of *δ*, except for *k*_on_, which is in units of *δ/*(1 c.u.). **(E)** Heatmap of the coupling score, *f*, with respect to *γ*_1,2_ and *γ*_2,3_, with the other parameters set to the most optimal parameter set (green curve in **D**), along with select choices of *γ*_1,2_ and *γ*_2,3_ (crosses) whose corresponding input-output curves are shown in **F. (F)** Input-output curves corresponding to the parameter sets indicated in **E. (G)** Overlap between the concentration ranges over which the input-output curves in **B** change by 90%. Given two intervals [*a, b*] and [*c, d*], the overlap is computed as max{0, min{*b, d*} *−* max{*a, c*}}*/*((*b − a*) + (*d − c*)). **(H)** Values of *γ*_1,2_ and *γ*_2,3_ for the best parameter set for each choice of RSC.

When examining the optimal parameter sets with a coupling score of *f <* 1 for RSC = 0.005, we found that *γ*_1,2_ and *γ*_2,3_ lie in the incoherent space, with *γ*_1,2_ *<* 1 and *γ*_2,3_ *>* 1 (Fig. 3C). To confirm the relevance of the incoherent regulatory mode, we identified the best parameter set from this ensemble of optimization results (Fig. 3D, green curve) and computed the coupling score while varying *γ*_1,2_ and *γ*_2,3_ within the range [10^−6^, 10^−6^](Fig. 3E shows a portion of this parameter space region; the green cross corresponds to the best parameter set). As expected, in this scenario, moving *γ*_1,2_ away from the identified minimum towards *γ*_1,2_ = 1 leads to a rapid increase in the coupling score, suggesting significant sensitivity of the coupling score to the value of *γ*_1,2_. To observe this more directly, we computed the responses for the parameter points corresponding to the crosses in Fig. 3E. When we set *γ*_1,2_ = 1 (black cross), we obtained 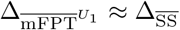, indicating significantly reduced decoupling; and when we entered the coherent space by setting *γ*_1,2_ *>* 1 (pink cross), we found that 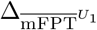 exceeds 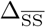 (Fig. 3F).

By analysing the curves in Fig. 3B, we found that the concentration range over which the steady-state level changed the most was systematically different from that for the mFPT, which can be quantified by the overlap of the input ranges over which the curves change the most (Fig. 3G, see also the green curves in Fig. 3F). Examining the parameter sets, we noticed that the concentration, *x*_1*/*2_, at which the steady-state level is half-maximal, i.e.,

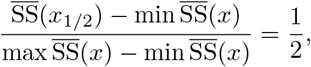

is always close to *k*_off_*/k*_on_, which is reminiscent of a Michaelis–Menten kinetic scheme (Fig. S3A, left). We also observed that the concentration at which the normalised mFPT is minimised, which we denote by *x*_fast_, is close to *ℓ*_1,2_*/k*_on_ (Fig. S3A, right). Together, this suggests that we can modulate the overlap to some extent by tuning *k*_off_. Indeed, we found that increasing the value of *k*_off_ in the best parameter set in Fig. 3D shifted the normalised steady-state curve rightward and increased *x*_1*/*2_, while only minimally affecting *x*_fast_ (Fig. S3B). However, we also found that increasing *k*_off_ beyond a certain critical value also increases 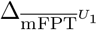 (Fig. S3B and C). This illustrates that, within an appropriate range of values of *k*_off_, we not only achieve global decoupling in the sense that the variation in the steady-state is much larger than that of the activation time, but we can also achieve a concentration-dependent form of decoupling, in which 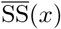 and 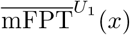 both vary with *x*, but over largely non-overlapping ranges.

Regarding the effect of the *RSC* value and thus the similarity of the rates of the various transitions, we found that, as we increased RSC to allow for rate scale separation (RSC ≥ 1), the optimal value of *γ*_1,2_ approached 1, whereas the optimal value of *γ*_2,3_ remained similarly large (Fig. 3H). In other words, we observed a transition from an incoherent regime, in which *γ*_1,2_ *<* 1, *γ*_2,3_ ≫ 1, and the transition rates are more tightly constrained, to a regime in which *B*_1_ → *B*_2_ is unregulated (*γ*_1,2_ ≈ 1), *γ*_2,3_ ≫ 1, and the transition rates are separated (i.e., case 1.III in Table 1, Fig. 2E).

We also noticed that the optimisations tended to yield values of *r* near the maximum possible value (*r* ≈ 10^4^, Fig. 3D), although there was a spread of values. To ascertain whether a large value of *r* is necessary for decoupling in this context, we also ran optimisations with the additional constraints that *ℓ*_1,2_ = *ℓ*_2,3_ = *r* and *ℓ*_2,1_ = *ℓ*_3,2_, so that the production transition proceeds on the same timescale as the preceding forward transitions, *U*_1_ *→ U*_2_ and *U*_2_ → *U*_3_ (Fig. S4). As before, we observed the strongest decoupling when the ligand operates in an incoherent regime, with comparably low coupling scores as in the previous optimisation (Fig. S4). This suggests that decoupling due to incoherent regulation does not require a large value of *r* relative to the other transition rates.

Finally, we also considered regulatory regimes in which one or both of the backward transitions, *B*_2_ *→ B*_1_ and *B*_3_ *→ B*_2_, are regulated. Specifically, we performed additional optimisations with the constraints *ℓ*_1,2_ = *ℓ*_2,3_ = *r* and *ℓ*_2,1_ = *ℓ*_3,2_, but with the following choices of regulatory factors that may differ from 1: (I) *γ*_2,1_ and/or *γ*_2,3_; (II) *γ*_1,2_ and/or *γ*_3,2_; or (III) *γ*_2,1_ and/or *γ*_3,2_ (Figs. S6, S7, and S8, respectively; Supporting Information S4). Here, we found instances of decoupling for cases II and III in the incoherent space (*γ*_1,2_ *<* 1 and *γ*_3,2_ *<* 1 for case II, *γ*_2,1_ *>* 1 and *γ*_3,2_ *<* 1 for case III), in which the ligand-bound transitions were biased towards *B*_1_ and *B*_3_. This again demonstrates that incoherent regulation can give rise to decoupling. On the other hand, no instances of decoupling were found for case I, for which the reasons remain elusive (Fig. S6).

In summary, our findings demonstrate that, when the transition rates are similar to each other, incoherent regulation can facilitate decoupling between the steady-state level and activation time. In this regime, unlike decoupling due to rate scale separation, the activation time exhibits a more significant dependence on the input concentration, but this variation can occur over a concentration range that is largely non-overlapping with that over which the steady-state changes (Fig. 3G), effectively leading to decoupling. We have also confirmed that these results extend to the larger graph, 𝒟 _6_ (Fig. S9D-E).

### Decoupling from an equilibrium of initial states

So far, we have defined the activation time in the ladder models, 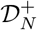, as the mFPT from one initial state, *U*_1_, to the terminal state, *M*, in which a copy of the readout *M* has been produced. This definition is reasonable in the setting of morphogen-mediated gene regulation in developmental systems such as the *Drosophila* blastoderm, a key model system that motivated our analyses in this paper [9, 11, 14, 16]. In this system, the nuclei divide every few minutes, with each mitosis resulting in the repression of transcription and the condensation of chromatin into a broadly inaccessible state. Therefore, it is reasonable in this context to assume that, upon the initiation of a new nuclear cycle, the regulatory DNA element that binds the morphogen has been “reset” to exist in the state *U*_1_.

However, in other contexts, such as those in which the cell is non-dividing or exhibits a long division time, it is plausible that the system exists in an equilibrium of initial states before the ligand is introduced. In this case, a more appropriate measure of the activation time would be the average mFPT to the terminal state over all possible initial states, each weighted by its steady-state probability (Fig. 4A). In other words, we define,

**Fig. 4.**
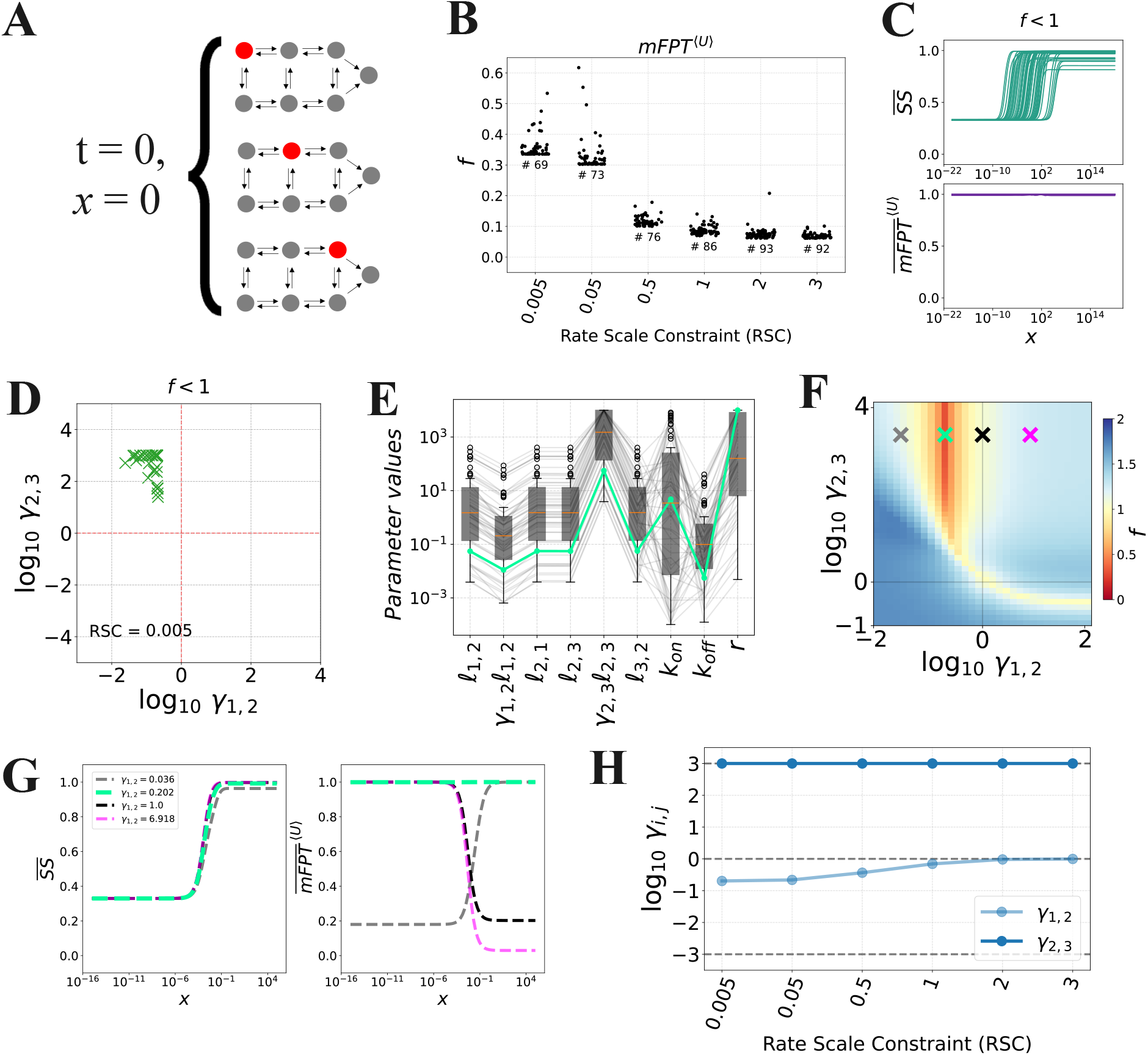
Decoupling from an equilibrium of initial states. **(A)** Schematic of the mFPT^*⟨U⟩*^(*x*) definition of activation time. This definition assumes that, before the ligand is introduced, the system has reached a steady state over the unbound states. **(B)** Distributions of the coupling score, *f*, obtained from optimization with various rate scale constraints (RSC). Input-output responses corresponding to the parameter sets obtained from optimization with RSC = 0.005. **(D)** Values of *γ*_1,2_ and *γ*_2,3_ for optimal parameter sets for RSC = 0.005. **(E)** Values of the parameter sets from each optimization run for RSC = 0.005. The green line represents the best parameter set. Each parameter is plotted in units of *δ*, except for *k*_on_, which is in units of *δ/*(1 c.u.). **(F)** Heatmap of the coupling score, *f*, with respect to *γ*_1,2_ and *γ*_2,3_, with the other parameters set to the most optimal parameter set (green curve in **E**), along with select choices of *γ*_1,2_ and *γ*_2,3_ (crosses) whose corresponding input-output curves are shown in **G. (G)** Input-output curves corresponding to the parameter sets indicated in **F. (H)** Values of *γ*_1,2_ and *γ*_2,3_ for the best parameter set for each choice of RSC.

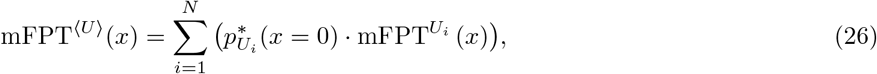

where we have used the same notation as in Eqn. 4, and 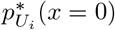 is the steady-state probability of *U*_*i*_ in 𝒟_*N*_ prior to introduction of ligand (*x* = 0). A mathematical justification of this definition is provided in Supporting Information S5.

To determine whether decoupling can be obtained with this alternative definition of activation time, we ran the constrained PSO with different choices of RSC (Eqn. 25), again focusing on the case where *γ*_1,2_ and *γ*_2,3_ may be distinct from 1, and *γ*_2,1_ = *γ*_3,2_ = 1. As with the preceding analysis, we found that the strength of decoupling decreases as we decrease the RSC (corresponding to a stronger constraint on the transition rates), but we still observed significant decoupling at RSC = 0.005 (Fig. 4B). As before, the increase in the coupling score with the RSC can be attributed to a smaller dynamic range in the steady-state level (Fig. 4C, top). In this case, we found that the activation time shows little to no dependence on the ligand concentration, in contrast to our previous analysis (Fig. 4C, bottom; compare to Fig. 3B, bottom). We confirmed this observation with Gillespie simulations, which we performed to further validate the definition in Eqn. 26 (Fig. S5).

Importantly, the optimized parameter sets were found to lie in the incoherent space for RSC = 0.005 (Fig. 4D). Upon varying the regulatory factors, *γ*_1,2_ and *γ*_2,3_, in the best parameter set (Fig. 4E, green curve), we again found that the strongest decoupling is indeed obtained when *γ*_1,2_ *<* 1 and *γ*_2,3_ *>* 1 (Figs. 4F and G). Moreover, increasing RSC results in optimal parameter sets that satisfy *γ*_1,2_ ≈ 1 and *γ*_2,3_ *>* 1 (Fig. 4H), again demonstrating that, as we allow for rate scale separation, we reach regulatory regimes in which only *B*_2_ *→ B*_3_ is regulated (case 1.III in Table 1). Finally, we confirmed that decoupling under incoherent regulation with respect to 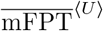 can also be observed in the 𝒟 _6_ model (Fig. S9F–G).

## Discussion

In this work, we sought to identify regulatory strategies that enable an input ligand to modulate the steady-state level of a molecular readout while maintaining a constant activation time, i.e., the time for the first readout molecule to be produced. By systematically analyzing Markov process models of an input-output system in which a ligand binds to a single regulatory site, we found two distinct regimes that support such decoupling between steady-state level and activation time. In the first regime, which we call *rate scale separation*, different transitions in the system proceed on different timescales, the system first undergoes slow transitions and then fast transitions, and the ligand does not regulate the slow forward transitions. In this way, the ligand controls the steady-state level whereas the slow, rate-limiting forward transitions dictate the activation time in a ligand-independent manner. In the second regime, the ligand acts as an *incoherent regulator*, exerting mutually opposing effects on different transitions towards the productive state.

The rate scale separation scenario aligns well with the setting of transcriptional regulation, where it is well-known that different regulatory steps—such as chromatin remodeling, transcription initiation, and RNA polymerase pausing—operate on distinct timescales (reviewed in [41]), and TFs may selectively regulate a subset of these processes [26, 42, 43]. In this way, a TF may regulate the steady-state level without affecting the activation time.

This idea is in line with the data reported in Eck *et al*. [9] on the regulation of *hunchback* by Bicoid and Zelda in the *Drosophila* blastoderm. In that paper, the authors report that in WT embryos, a *hunchback* reporter exhibits a constant activation time over the antero-posterior axis of the embryo, despite the presence of a Bicoid concentration gradient along this axis, which only affects the reporter transcription levels. However, in embryos lacking the pioneer factor Zelda, which is regarded to be present at a roughly constant level along the antero-posterior axis, both the activation time and the transcription level were found to depend on Bicoid concentration. One plausible interpretation of this data, according to our findings here, is that the activation time is determined by the activity of Zelda, which promotes chromatin opening constantly throughout the embryo, which is putatively a slow process, whereas the RNA polymerase loading rate, which corresponds to transcription levels, proceeds on a faster timescale and is regulated by Bicoid. In Supporting Information S6 and Fig. S11 we show an adaptation of the 𝒟 _3_ model to qualitatively account for this data.

In a more recent paper, Alamos *et al*. [25] reported similar observations of decoupling with a synthetic enhancer that binds the morphogen Dorsal in the *Drosophila* embryo. However, Alamos *et al*. proposed a different model, namely a model based on the Erlang process [44], to explain this data. This model is equivalent to a version of the chain model, 𝒞 _*N*_, in which all backward transitions are omitted and all forward transitions have the same rate, which depends on Dorsal concentration. While Alamos *et al*. showed that this model can recapitulate the observed decoupling on average, their analysis also assumed a short time window (∼ 7 min) during which transcription could be activated. Such a time window is naturally imposed by the fast nuclear division times in the early *Drosophila* embryo [45], but this assumption also effectively truncates the distribution of possible activation times in a fashion that causes the mean of this distribution to appear independent of Dorsal concentration (Fig. S12A,D). However, we have found that allowing for longer observation windows in the same model yields an activation time that *does* depend on input concentration (Supporting Information S7, Fig. S12D). In light of this observation, the regulatory strategies we have proposed in this paper can serve as alternative explanations of the decoupling observed by Alamos *et al*.. This emphasises the importance of considering alternative models when interpreting experimental data, and showcases how this can also inform experimental design and interpretation, in this case highlighting the possible limitations imposed by short measurement windows, which have also been noted previously [45].

In addition to rate scale separation, we have also found that decoupled responses can arise under incoherent regulation. Such regulatory regime is increasingly recognized as a feature of gene regulation [23, 46], but a thorough mechanistic understanding of how incoherence arises, and what functional consequences this regulatory mode has, is still lacking. Our analysis reveals that one such consequence may be decoupling, albeit to a less robust extent than the decoupling obtained under rate scale separation. Our optimization results in Figs. 3 and S6–S8 also suggest that the ability of an incoherent regulator to achieve decoupling may depend on which transitions are regulated. We envision that further work should aim to identify the origins of these dependencies, as well as the differing constraints and tradeoffs that apply to systems under rate scale separation and incoherent regulation.

Among the incoherent parameter sets that exhibit decoupling with respect to the 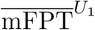 definition of activation time, we noted that those pertaining to some regulatory regimes have low unbinding rates, *k*_off_ (Figs. 3D and S7). However, in the context of gene regulation, experimental measurements of TF–DNA dissociation rates generally tend to be fast [40, 41]. This is more consistent with the results in Fig. S8, which show that when *B*_2_ *→ B*_1_ and *B*_3_ *→ B*_2_ are regulated, larger values of *k*_off_ can support decoupling. In addition, our analysis of decoupling with respect to 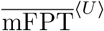 yielded a collection of parameter sets with greater variability in *k*_off_, and we were indeed able to identify parameter sets with larger *k*_off_ values that support decoupling. We suspect that the need for a relatively low *k*_off_ in some regulatory regimes arises from our assumption that regulation occurs only while the ligand is bound. We anticipate that decoupling and fast unbinding can co-occur more consistently in more complex models in which the regulatory effect of the ligand persists even after unbinding, through the inclusion of “memory states.”

Our analysis focused on particular choices of models—the chain models, 𝒞_*N*_, and the ladder models, 𝒟_*N*_ —alongside additional assumptions, such as the assumption that the ligand regulates at most two horizontal transitions in 𝒟_*N*_ . As such, there are several aspects of this analysis that may be generalized in future work. First, examining the effects of different numbers of internal states (*N*), beyond the specific cases of *N* = 2, *N* = 3 and *N* = 6 considered here. Second, our models assume a single ligand-binding site, as is the case in many experimental systems [25, 26, 47, 48]; despite this, we also expect that incorporating multiple binding sites may yet reveal additional mechanisms for decoupling. In the same direction, incorporating multiple ligands would be insightful. Third, in the ladder models, we assumed that ligand unbinding proceeds with a constant rate, *k*_off_, that is independent of the system state. This assumption encodes the simple scenario in which progression towards the productive state does not influence the ligand’s unbinding kinetics. For example, a TF may bind to a regulatory element far from the promoter, but still affect the rate of polymerase assembly at the promoter in a way that does not modulate the TF binding properties. It is possible that additional regulatory strategies for decoupling may emerge upon relaxing this assumption.

We believe that the biological interpretation of the activation time as an mFPT in the underlying Markov process is straightforward. It is also experimentally tractable, as modern techniques now offer access to such timing information, often in the form of FPT distributions, in various molecular systems (e.g. [6, 8, 15, 49, 50]). On the other hand, other definitions of activation time may be considered. For example, recent studies in the literature [21, 51–54] have considered the time required for the molecular readout to reach a given mean abundance. Notably, this quantity depends on the degradation rate of the molecule, in contrast to our definition (Supporting Information S8, Fig. S10). It will be insightful to examine how decoupling arises for various definitions of activation time.

The relevance of the steady-state level as a measure of readout abundance is less clear, as there are many molecular systems that do not reach a steady state, especially in *in vivo* contexts in which the system is embedded in a highly dynamic environment. For instance, the transcriptional output of a gene rarely reaches a steady state, and RNA is typically produced in transient, stochastic bursts [55]. As such, while steady-state assumptions, or even assumptions of thermodynamic equilibrium, are widespread in theoretical studies of such systems [9, 27, 56–60], the overall validity of these assumptions are questionable. In any case, from a modelling perspective, checking for a large change in the steady state ensures that the flat activation timing is not a mere artifact of saturation or non-responsiveness of the system.

More broadly, decoupling between readout abundance and activation time has significant implications for input-output systems in biology. For example, cells across different spatial positions or lineages in a developmental system may need to respond synchronously to different local morphogen concentrations, which could be facilitated by a decoupling mechanism. Conversely, the ability to tune timing while maintaining fixed output levels could support functions that require fine-tuned activation timing regulation. These “reverse decoupling” regimes remain unexplored and merit further investigation. Additionally, applying similar analyses to alternative definitions of activation time—such as time to a threshold readout abundance [54], or to the onset of a burst in readout production—could deepen our understanding of dynamic regulatory mechanisms and improve the mechanistic interpretation of experimental data.

## Materials and Methods

### Calculating the steady-state level, activation time, and their dynamic ranges

As described in the section “Modeling approach and mathematical setup,” we used both numerical and analytical approaches for calculating the steady-state level, activation time, and their dynamic ranges, which we defined using the graph-theoretic linear framework [17–20] (Supporting Information S1). For the analytical calculations, we implemented a symbolic version of the Chebotarev–Agaev recurrence (Eqn. 10), using the Python package SymPy [61]. Barring additional symbolic simplifications, this approach allows us to obtain exact algebraic expressions for the steady-state level and activation time for any graph, *G*, in *O*(*n*^4^) arithmetic operations, where *n* is the number of vertices in *G*. This is much more efficient than any method based on direct enumeration of the spanning trees and/or forests that feature in Eqns. 6 and 7, which can scale exponentially with *n* [19].

For the numerical calculations, we calculated the steady-state level by obtaining the SVD of ℒ (*G*), identifying the right singular vector corresponding to the zero singular value, and normalizing appropriately. This right singular vector is unique whenever *G* is strongly connected. The normalized vector, which is the steady-state vector, **p**^*∗*^, arising from the master equation (Eqn. 1), was then used to evaluate 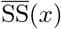, as per Eqns. 3 and 13. On the other hand, we calculated the activation time by obtaining the QR decomposition of the left-hand matrix in Eqn. 5, and using the corresponding solution vector to Eqn. 5 to evaluate 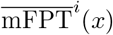, as per Eqn. 13. Briefly, a QR decomposition of an invertible matrix is a decomposition, **A** = **QR**, such that **Q** is orthogonal, **Q**^*−*1^ = **Q**^T^, and **R** is upper-triangular; such decompositions are useful for solving matrix equations of the form **Ax** = **b**. While the exact runtime complexities of these decompositions can differ between implementations, we expect that they will be asymptotically faster or comparable to the *O*(*n*^4^) runtime of the Chebotarev–Agaev recurrence, and—more importantly—faster in practice due to the availability of highly optimized implementations.

All calculations were implemented in C++, using multiple-precision floating-point numbers from the Boost.Multiprecision library [62] with a precision of 100 digits, and implementations of SVD and QR, from the Eigen library [63]. Python bindings were implemented to call these C++ functions from Python, so that they could interface with the PyMoo optimization suite (see below).

We checked that the mFPTs obtained from Eqn. 12 agree well with estimates obtained from Gillespie simulations [32, 64] on a simple four-vertex graph (Fig. S1A). We sampled 100 combinations of values for the rates, each from a log-uniform distribution on the range [10^*−*3^, 10^3^], and ran increasing numbers of Gillespie simulations starting from vertex 1 to estimate the mFPT to vertex 4. As expected, we observed that the agreement between these estimates and their corresponding exact values, given by Eqn. 12, increases with the number of trajectories (Fig. S1B).

To numerically compute the dynamic range for a given parameter set, we calculated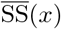 and 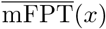 over a logarithmic concentration range of [10^*−*20^, 10^20^], with a logarithmic stepsize of ∼0.04004. We chose this wide range to ensure that these calculations capture the complete dynamic range of these outputs. For each output, we then identified the maximum and minimum value over this concentration range, and computed the dynamic ranges according to Eqn. 14.

### Minimization protocols for the decoupling score *f*

In preliminary investigations for this study, we compared various metaheuristic optimisation algorithms to find parameter sets that exhibit decoupling. We found the Particle Swarm Optimization (PSO) implementation provided by PyMoo [38], a multi-objective optimization suite, to be the most efficient option, and therefore used it for all the numerical analyses discussed in this work.

The PSO algorithm was first introduced by Kennedy and Eberhart in 1995 [37]. Conceptually, the procedure is based on a swarm of particles, each with an associated position and velocity vector, which are iteratively updated to minimise an objective function, *f* . We denote the position and velocity of the *i*-th particle along the *d*-th dimension at time *t* as 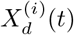 and 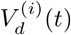, respectively. (Here, *t* is treated as an integer variable.) In our case, each particle corresponds to a parameter set, with the particle’s position given by the parameter values. Meanwhile, the velocity of the *i*-th particle along the *d*-th dimension, 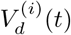, is determined by:

1. that particle’s position at which *f* attained its lowest value throughout the particle’s trajectory up to time *t*, which we denote by 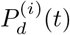; and
2. the position at which *f* attained its lowest value throughout the entire swarm’s history up to time *t*, which we denote by *G*_*d*_(*t*).

More specifically, 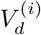 is given by

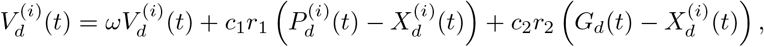

where *ω* is an inertia factor, *r*_1_, *r*_2_ *∈* [0, 1) are noise coefficients representing the level of “craziness” in the optimisation, and *c*_1_ and *c*_2_ balance the contributions from the *i*-th particle’s “personal” behaviour 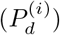 and the swarm’s global behaviour (*G*_*d*_) [37]. PyMoo dynamically adjusts *ω, c*_1_, and *c*_2_ throughout the optimisation, with initial values of *ω* = 0.9 and *c*_1_ = *c*_2_ = 2, following the prescription outlined in [65]. Finally, the position of each particle is updated as

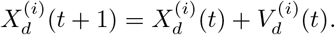

We ran PSO for each of the problems defined in the paper, passing a total of 100 different random seeds to obtain a population of optimal solutions. To sample the initial set of particles, we used Latin hypercube sampling, which is the default choice in PyMoo. To enforce parametric bounds, the parameters that fall outside the defined bounded range are set to the closest bound value during the optimisation.

As the termination criterion for each PSO run, we used either convergence to a score of *f <* 0.1 for more than five consecutive generations, or a computation time exceeding 23 hours. The termination criterion that was used for each PSO run is specified in the corresponding figure caption.

All optimisations were performed on the O2 High Performance Compute Cluster at Harvard Medical School.

## Code and data availability

All code and data used to generate the figures in this paper are available in GitHub at https://github.com/theobiolab/FPT_paper.git.

## Acknowledgments

The authors thank members of the Gunawardena group and Konstantina Poumpouridou for interesting discussions. This work was partially financed by NIH grant R01GM122928 (K-M.N. and J.G.), and grants from the Spanish Ministry of Science, Innovation and Universities MCIU/AEI/10.13039/501100011033 (grant RYC2021-033860-I cofounded by European Union NextGenerationEU/PRTR to R.M.-C.), and the Ministry of Science and Innovation (PID2022-142210NA-I00 funded by MCIN/AEI/10.13039/501100011033/FEDER, UE to R.M.-C.; and pre-doctoral fellowship PRE2022-101864 funded by MCIN/AEI/10.13039/501100011033 to G.R.) The authors acknowledge support of the Spanish Ministry of Science and Innovation through the Centro de Excelencia Severo Ochoa (CEX2020-001049-S, MCIN/AEI /10.13039/501100011033), the Generalitat de Catalunya through the CERCA programme and to the EMBL partnership. Research for this publication has been partially carried out in the Barcelona Collaboratorium for Modelling and Predictive Biology.

