## Supporting Information for "Decoupling between activation time and steady-state level in input-output responses"

#### S1 Derivations of steady-state level and activation time (Eqns. 3, 5, and 7)

As described in the section "Modeling approach and mathematical setup," we used the graph-theoretic linear framework [1–4] to formulate and analyze the models that we consider in this paper. Here, we provide brief mathematical derivations for our expressions for the steady-state level (Eqn. 3) and activation time (Eqns. 5 and 7). We will use the notation we have established above in what follows.

**Derivation of Eqn. 3.** First, we provide a short derivation of Eqn. 3, versions of which have also been provided in previous papers [5, 6]. To do this, we define an extended model that explicitly accounts for the copy-number of the molecular readout, $M$, produced from the system described by the graph, $G$, with productive vertices $\mathcal{V}_{\text{prod}}(G) \subset \mathcal{V}(G)$. As described above, this model describes the production of $M$ as proceeding at a constant rate, $r$, from each productive vertex. On the other hand, we assume that degradation occurs at a first-order rate, $\delta n_M$, that is independent of the system state. The corresponding Markov process may be viewed as arising from an infinite "copy-number graph" [6], whose vertices, $(i, n_M) \in \mathcal{V}(G) \times \mathbb{R}_{\geq 0}$, keep track of both the system state, $i \in \mathcal{V}(G)$, and the readout copy-number, $n_M$. The corresponding master equation is given by

$$\frac{dp_{i,n_M}(t)}{dt} = r_i p_{i,n_M-1}(t) + (n_M+1)\,\delta p_{i,n_M+1}(t) - (r_i + \delta n_M)\,p_{i,n_M}(t) + \sum_{j\in\mathcal{V}(G)} \mathcal{L}(G)_{i,j}\,p_{j,n_M}(t),$$

where we have defined

$$r_i = \begin{cases} r & \text{if } i \in \mathcal{V}_{\text{prod}}(G) \\ 0 & \text{otherwise,} \end{cases}$$

and we appropriately omit the first right-hand term when $n_M = 0$. Now, let us assume that $\mathcal{V}(G) = \{1, \ldots, n\}$, and define the vector, $\mathbf{p}_{n_M}(t) = (p_{1,n_M}(t), \ldots, p_{N,n_M}(t))^{\mathrm{T}}$. Then the above master equation implies a corresponding master equation for $\mathbf{p}_{n_M}(t)$, as

$$\frac{d}{dt}\mathbf{p}_{n_M}(t) = \mathbf{R}\mathbf{p}_{n_M-1}(t) + (n_M+1)\,\delta\mathbf{p}_{n_M+1}(t) - (\mathbf{R} + \delta n_M \mathbf{I} - \mathcal{L}(G))\,\mathbf{p}_{n_M}(t), \tag{S1}$$

where $\mathbf{R}$ is the $n \times n$ diagonal matrix with entries $R_{i,i} = r_i$. We now wish to derive an expression for the mean steady-state value of $n_M$, which is given by

$$\langle n_M \rangle^* = \sum_{n_M=1}^{\infty} n_M \sum_{i \in \mathcal{V}(G)} p^*_{i,n_M} = \mathbf{1}^{\mathrm{T}} \boldsymbol{\mu}^*, \qquad \text{where} \qquad \boldsymbol{\mu}^* = \sum_{n_M=1}^{\infty} n_M \mathbf{p}^*_{n_M},$$

and $\mathbf{1}$ is the all-ones vector of dimension $N$.

First, let $q_i(t)$ be the marginal probability of the system state $i \in \mathcal{V}(G)$ at time $t$,

$$q_i(t) = \sum_{n_M=0}^{\infty} p_{i,n_M}(t),$$

and let $\mathbf{q}(t) = (q_1(t), \dots, q_N(t))^{\mathrm{T}}$. Then Eqn. S1 tells us that

$$\begin{aligned} \frac{d}{dt}\mathbf{q}(t) &= \sum_{n_M=0}^{\infty} \frac{d}{dt}\mathbf{p}_{n_M}(t) \\ &= \underbrace{\delta \mathbf{p}_1(t) - \mathbf{R}\mathbf{p}_0(t) + \mathcal{L}(G)\,\mathbf{p}_0(t)}_{n_M=0} + \underbrace{\mathbf{R}\mathbf{p}_0(t) + 2\delta\mathbf{p}_2(t) - \mathbf{R}\mathbf{p}_1(t) - \delta\mathbf{p}_1(t) + \mathcal{L}(G)\,\mathbf{p}_1(t)}_{n_M=1} \\ &\quad + \underbrace{\mathbf{R}\mathbf{p}_1(t) + 3\delta\mathbf{p}_3(t) - \mathbf{R}\mathbf{p}_2(t) - 2\delta\mathbf{p}_2(t) + \mathcal{L}(G)\,\mathbf{p}_2(t)}_{n_M=2} + \cdots, \end{aligned}$$

which can be easily rearranged as

$$\frac{d}{dt}\mathbf{q}(t) = \mathcal{L}(G)\left(\mathbf{p}_0(t) + \mathbf{p}_1(t) + \mathbf{p}_2(t) + \cdots\right) = \mathcal{L}(G)\,\mathbf{q}(t).$$

Therefore, the steady-state marginal probability vector, $\mathbf{q}^*$, must lie in $\ker \mathcal{L}(G)$, and must in fact equal the vector, $\mathbf{p}^*$, of steady-state probabilities in $G$ (Eqns. 3 and 6).

We are now ready to derive a master equation for the vector,

$$\boldsymbol{\mu}(t) = \sum_{n_M=1}^{\infty} n_M \mathbf{p}_{n_M}(t),$$

whose steady state was defined above. Applying Eqn. S1, we can write

$$\begin{aligned} \frac{d}{dt}\boldsymbol{\mu}(t) &= \sum_{n_M=1}^{\infty} n_M \frac{d}{dt}\mathbf{p}_{n_M}(t) \\ &= \sum_{n_M=1}^{\infty} n_M \left(\mathbf{R}\mathbf{p}_{n_M-1}(t) + (n_M+1)\,\delta\mathbf{p}_{n_M+1}(t) - (\mathbf{R} + \delta n_M \mathbf{I} - \mathcal{L}(G))\,\mathbf{p}_{n_M}(t)\right), \end{aligned}$$

which can be reorganized as the sum of three terms,

$$\begin{aligned} \frac{d}{dt}\boldsymbol{\mu}(t) = \sum_{n_M=1}^{\infty} n_M \mathbf{R}\left(\mathbf{p}_{n_M-1}(t) - \mathbf{p}_{n_M}(t)\right) + \sum_{n_M=1}^{\infty} \delta n_M \left((n_M+1)\,\mathbf{p}_{n_M+1}(t) - n_M \mathbf{p}_{n_M}(t)\right) \\ + \sum_{n_M=1}^{\infty} n_M \mathcal{L}(G)\,\mathbf{p}_{n_M}(t). \end{aligned} \tag{S2}$$

We now consider each of the three terms. The first simplifies to

$$\begin{aligned} \sum_{n_M=1}^{\infty} n_M \mathbf{R}\left(\mathbf{p}_{n_M-1}(t) - \mathbf{p}_{n_M}(t)\right) &= \mathbf{R}\left(\mathbf{p}_0(t) - \mathbf{p}_1(t) + 2\mathbf{p}_1(t) - 2\mathbf{p}_2(t) + 3\mathbf{p}_2(t) - + \cdots\right) \\ &= \mathbf{R}\mathbf{q}(t). \end{aligned}$$

The second term simplifies to

$$\begin{aligned}\sum_{n_M=1}^{\infty} &\delta n_M \left(\left(n_M+1\right)\mathbf{p}_{n_M+1}(t) - n_M \mathbf{p}_{n_M}(t)\right) \\ &= \delta\left(\left(2\mathbf{p}_2(t) - \mathbf{p}_1(t)\right) + 2\left(3\mathbf{p}_3(t) - 2\mathbf{p}_2(t)\right) + \cdots\right) - \delta\left(\mathbf{p}_1(t) + 2\mathbf{p}_2(t) + \cdots\right) \\ &= -\delta\boldsymbol{\mu}(t).\end{aligned}$$

Finally, the third term is simply

$$\sum_{n_M=1}^{\infty} n_M \mathcal{L}(G)\,\mathbf{p}_{n_M}(t) = \mathcal{L}(G)\,\boldsymbol{\mu}(t).$$

As such, we can rewrite Eqn. S2 as simply

$$\frac{d}{dt}\boldsymbol{\mu}(t) = \mathbf{R}\mathbf{q}(t) - \delta\boldsymbol{\mu}(t) + \mathcal{L}(G)\,\boldsymbol{\mu}(t), \tag{S3}$$

which, at steady state, becomes

$$\delta\boldsymbol{\mu}^* - \mathcal{L}(G)\,\boldsymbol{\mu}^* = \mathbf{R}\mathbf{q}^*.$$

Now, we left-multiply both sides by $\mathbf{1}^{\mathrm{T}}$, to get

$$\delta\mathbf{1}^{\mathrm{T}}\boldsymbol{\mu}^* - \mathbf{1}^{\mathrm{T}}\mathcal{L}(G)\,\boldsymbol{\mu}^* = \mathbf{1}^{\mathrm{T}}\mathbf{R}\mathbf{q}^*.$$

It follows from the definition of $\mathcal{L}(G)$ (Eqn. 2) that the columns of $\mathcal{L}(G)$ sum to zero. Therefore, $\mathbf{1}^{\mathrm{T}}\mathcal{L}(G)$ is the zero vector, and we get

$$\langle n_M\rangle^* = \mathbf{1}^{\mathrm{T}}\boldsymbol{\mu}^* = \frac{\mathbf{1}^{\mathrm{T}}\mathbf{R}\mathbf{q}^*}{\delta}.$$

Now, recalling the definition of $\mathbf{R}$ and the fact that $\mathbf{q}^* = \mathbf{p}^*$, we finally obtain

$$\langle n_M\rangle^* = \frac{1}{\delta}\sum_{i\in\mathcal{V}(G)} r_i p_i^* = \frac{r}{\delta}\sum_{i\in\mathcal{V}_{\mathrm{prod}}(G)} p_i^*,$$

i.e., we recover Eqn. 3.

**Derivation of Eqn. 5.** We now turn to a derivation of Eqn. 5, which has also been provided in previous work [7]. Recalling the notation introduced above, let $G$ be a strongly connected graph that describes the input-output system of interest, with vertices $\mathcal{V}(G) = \{1,\ldots,n\}$, of which $\mathcal{V}_{\mathrm{prod}}(G) \subset \mathcal{V}(G)$ is the subset of productive vertices; and let $G^+$ be the graph obtained by adding a new vertex, $M$, together with edges $j \to M$ for each $j \in \mathcal{V}_{\mathrm{prod}}(G)$. We defined the activation time as the mFPT from a chosen initial vertex, $i \in \mathcal{V}(G)$, to $M$ in the corresponding Markov process, $X^+(t)$, which can be written as

$$\mathrm{mFPT}^i = \mathbb{E}\left[\inf\left\{t > 0 : X^+(t) = M\right\} \mid X^+(0) = i\right].$$

Now, let $w_i(t)$ denote the probability that a trajectory of $X^+(t)$ that begins at $i$ reaches $M$ by time $t$,

$$w_i(t) = \Pr\left[X^+(t') = M \text{ for some } 0 \le t' \le t \mid X^+(0) = i\right].$$

Since $G$ is strongly connected, any such trajectory of $X^+(t)$ will eventually reach $M$ with probability one, i.e., $\lim_{t\to 0} w_i(t) = 1$. Now, it can be shown [7] that both $w_i(t)$ and its density, which we denote by $u_i(t) = dw_i/dt$, each satisfies an *adjoint master equation*,

$$\frac{d}{dt}\mathbf{w}(t) = \left(\mathcal{L}(G^+)_{[\overline{\{n+1\}},\overline{\{n+1\}}]}\right)^{\mathrm{T}}\mathbf{w}(t) \tag{S4}$$

$$\frac{d}{dt}\mathbf{u}(t) = \left(\mathcal{L}(G^+)_{[\overline{\{n+1\}},\overline{\{n+1\}}]}\right)^{\mathrm{T}}\mathbf{u}(t). \tag{S5}$$

Here, we have defined the vectors $\mathbf{w}(t) = (w_1(t), \ldots, w_n(t))^{\mathrm{T}}$ and $\mathbf{u}(t) = (u_1(t), \ldots, u_n(t))^{\mathrm{T}}$, we have identified $M \equiv n+1$, and $\mathcal{L}(G^+)_{[\overline{\{n+1\}},\overline{\{n+1\}}]}$ is the submatrix of $\mathcal{L}(G^+)$ obtained by removing the row and column corresponding to $M \equiv n+1$.

Now, let $\tilde{u}_i(s)$ be the Laplace transform of $u_i(t)$,

$$\tilde{u}_i(s) = \int_0^\infty e^{-st} u_i(t)\, dt.$$

Taking Laplace transforms of both sides of Eqn. S5, we get

$$s\tilde{\mathbf{u}}(s) - \mathbf{u}(0) = \left(\mathcal{L}(G^+)_{[\overline{\{n+1\}},\overline{\{n+1\}}]}\right)^{\mathrm{T}} \tilde{\mathbf{u}}(s), \tag{S6}$$

where $\tilde{\mathbf{u}}(s) = (\tilde{u}_1(s), \ldots, \tilde{u}_n(s))^{\mathrm{T}}$. To determine the entries in $\mathbf{u}(0)$, we can turn to Eqn. S4, which tells us that

$$u_i(0) = \left.\frac{dw_i}{dt}\right|_{t=0} = \sum_{j \in \mathcal{V}(G)} \ell(i \to j)\, w_j(0) - \left(\sum_{j \in \mathcal{V}(G^+)} \ell(i \to j)\right) w_i(0).$$

Since $w_M(0) = 1$ and $w_j(0) = 0$ for all $j \neq M$, this implies that

$$u_i(0) = \begin{cases} \ell(i \to M) & \text{if } i \in \mathcal{V}_{\mathrm{prod}}(G) \\ 0 & \text{otherwise.} \end{cases}$$

Hence, Eqn. S6 can be rearranged as

$$\left(\mathbf{L}(G^+)_{[\overline{\{n+1\}},\overline{\{n+1\}}]} + s\mathbf{I}\right) \tilde{\mathbf{u}}(s) = \begin{bmatrix} \ell(1 \to M) \\ \vdots \\ \ell(n \to M) \end{bmatrix},$$

or

$$\tilde{\mathbf{u}}(s) = \left(\mathbf{L}(G^+)_{[\overline{\{n+1\}},\overline{\{n+1\}}]} + s\mathbf{I}\right)^{-1} \begin{bmatrix} \ell(1 \to M) \\ \vdots \\ \ell(n \to M) \end{bmatrix}, \tag{S7}$$

where we have defined $\mathbf{L}(G^+) = -\mathcal{L}(G^+)^{\mathrm{T}}$ and we have implicitly set $\ell(i \to M) \equiv 0$ if the edge does not exist.

Now, let $\mu_i^{(r)}$ be the $r$-th moment of $u_i(t)$,

$$\mu_i^{(r)} = \mathbb{E}\left[\inf\left\{t > 0 : X^+(t) = M\right\}^r \mid X^+(0) = i\right] = \int_0^\infty t^r u_i(t)\, dt.$$

We utilize the simple fact that the moments of $u_i(t)$ are related to $\tilde{u}_i(s)$, as

$$\mu_i^{(r)} = \int_0^\infty t^r u_i(t)\, dt = (-1)^r \frac{d^r}{ds^r} \int_0^\infty e^{-st} u_i(t)\, dt\Big|_{s=0} = (-1)^r \frac{d^r \tilde{u}_i}{ds^r}\Big|_{s=0}.$$

Therefore, if we define the vector of moments, $\boldsymbol{\mu}^{(r)} = \left(\mu_1^{(r)}, \ldots, \mu_n^{(r)}\right)^{\mathrm{T}}$, we can rewrite Eqn. S7 as

$$\boldsymbol{\mu}^{(r)} = (-1)^r \frac{d^r}{ds^r} \left(\mathbf{L}(G^+)_{[\overline{\{n+1\}},\overline{\{n+1\}}]} + s\mathbf{I}\right)^{-1}\Big|_{s=0} \begin{bmatrix} \ell(1 \to M) \\ \vdots \\ \ell(n \to M) \end{bmatrix},$$

which can be written as [8]

$$\boldsymbol{\mu}^{(r)} = r! \left(\mathbf{L}(G^+)_{[\overline{\{n+1\}},\overline{\{n+1\}}]}\right)^{-(r+1)} \begin{bmatrix} \ell(1 \to M) \\ \vdots \\ \ell(n \to M) \end{bmatrix}.$$

Setting $r = 1$, we finally recover Eqn. 5.

**Derivation of Eqn. 7.** Finally, we provide a brief derivation of Eqn. 7, again leaving the details to a previous paper [7]. To do this, we turn to the *All-Minors Matrix-Tree theorem* (AMMTT), which relates the *minors* (determinants of submatrices) of $\mathcal{L}(G)$ to the spanning forests of $G$ [9]. It is through this fundamental relationship that the spanning forests of $G$ determine both $\mathrm{SS}(x)$ (Eqns. 3 and 6) and $\mathrm{mFPT}^i(x)$ (Eqn. 7); see [3, 7] for further details.

Let $G$ be a strongly connected graph on vertices $\mathcal{V}(G) = \{1, \ldots, n\}$, and let $G^+$ be the graph obtained by adding a new vertex, $M \equiv n+1$, as described in the main text. Then the AMMTT tells us that, for any $i, j \in \mathcal{V}(G)$, we have [7]

$$\det \mathcal{L}(G^+)_{[\overline{\{i,n+1\}},\overline{\{j,n+1\}}]} = (-1)^{n-1+i+j}\, w(\Phi_{\{j,n+1\}:i\rightsquigarrow j}(G^+)), \tag{S8}$$

where we have used the weight function, $w(\cdot)$, introduced in the main text; and, in analogy with the notation introduced in Eqn. 5, $\mathcal{L}(G^+)_{[\overline{\{i,n+1\}},\overline{\{j,n+1\}}]}$ is the submatrix of $\mathcal{L}(G^+)$ obtained by removing the rows indexed by $\{i, n+1\}$ and the columns indexed by $\{j, n+1\}$. Meanwhile, the AMMTT also tells us that [2]

$$\det \mathcal{L}(G^+)_{[\overline{\{n+1\}},\overline{\{n+1\}}]} = (-1)^{n}\, w(\Phi_{\{n+1\}}(G^+)). \tag{S9}$$

Now, let us recall Cramer's rule, which states that the inverse of any invertible matrix, $\mathbf{A}$, is given by

$$\mathbf{A}^{-1} = \frac{1}{\det \mathbf{A}} \left(\operatorname{adj} \mathbf{A}\right),$$

where $\operatorname{adj}\mathbf{A}$ is the adjugate matrix of $\mathbf{A}$, whose $(i,j)$-th entry is given by

$$\left(\operatorname{adj} \mathbf{A}\right)_{i,j} = (-1)^{i+j} \det \mathbf{A}_{[\overline{\{j\}},\overline{\{i\}}]}.$$

Combining this with Eqns. S8 and S9, we can now evaluate the $(i,j)$-th entry of the matrix $\left(\mathbf{L}(G^+)_{[\overline{\{n+1\}},\overline{\{n+1\}}]}\right)^{-1}$, as

$$\begin{aligned}
\left(\mathbf{L}(G^+)_{[\overline{\{n+1\}},\overline{\{n+1\}}]}\right)^{-1}_{i,j} &= -\left(\mathcal{L}(G^+)_{[\overline{\{n+1\}},\overline{\{n+1\}}]}\right)^{-1}_{j,i} \\
&= -\left(\frac{\operatorname{adj}\left(\mathcal{L}(G^+)_{[\overline{\{n+1\}},\overline{\{n+1\}}]}\right)}{\det \mathcal{L}(G^+)_{[\overline{\{n+1\}},\overline{\{n+1\}}]}}\right)_{j,i} \\
&= -\frac{(-1)^{i+j} \det \mathcal{L}(G^+)_{[\overline{\{i,n+1\}},\overline{\{j,n+1\}}]}}{\det \mathcal{L}(G^+)_{[\overline{\{n+1\}},\overline{\{n+1\}}]}} \\
&= -\frac{(-1)^{n-1+2i+2j}\, w(\Phi_{\{j,n+1\}:i\rightsquigarrow j}(G^+))}{(-1)^{n}\, w(\Phi_{\{n+1\}}(G^+))} \\
&= \frac{w(\Phi_{\{j,n+1\}:i\rightsquigarrow j}(G^+))}{w(\Phi_{\{n+1\}}(G^+))}.
\end{aligned}$$

Now, we combine this with Eqn. 5, to finally obtain

$$\begin{aligned}
\mathrm{mFPT}^i(x) &= \sum_{j=1}^{n} \left(\mathbf{L}(G^+)_{[\overline{\{n+1\}},\overline{\{n+1\}}]}\right)^{-2}_{i,j} \ell(j \to n+1) \\
&= \sum_{j=1}^{n} \left(\mathbf{L}(G^+)_{[\overline{\{n+1\}},\overline{\{n+1\}}]}\right)^{-1}_{i,j} \sum_{k=1}^{n} \left(\mathbf{L}(G^+)_{[\overline{\{n+1\}},\overline{\{n+1\}}]}\right)^{-1}_{j,k} \ell(k \to n+1) \\
&= \sum_{j=1}^{n} \frac{w(\Phi_{\{j,n+1\}:i\rightsquigarrow j}(G^+))}{w(\Phi_{\{n+1\}}(G^+))} \sum_{k=1}^{n} \left(\frac{w(\Phi_{\{k,n+1\}:j\rightsquigarrow k}(G^+))}{w(\Phi_{\{n+1\}}(G^+))}\right) \ell(k \to n+1).
\end{aligned}$$

It is easy to show, using an elementary graph-theoretic argument, that the inner sum evaluates to 1 [7]. This brings us to Eqn. 7.

#### S2 Monotonicity of steady-state level and activation time with input concentration under single-rate or coherent regulation

In this section, we outline our strategy for calculating the sign of the derivative of the steady-state level, $\mathrm{SS}(x)$, and activation time, $\mathrm{mFPT}^{U_1}(x)$, with respect to the ligand concentration, $x$, in the ladder models $\mathcal{D}_2$ and $\mathcal{D}_3$. This will allow us to show that, under certain assumptions, the two outputs are monotonic in $x$. Due to the complexity of the underlying equations, the analytics is mostly based on symbolic computation using Mathematica and SymPy [10].

In particular, we shall show that, for $\mathcal{D}_2$,

$$\begin{aligned}
\gamma_{1,2} > 1 \quad \text{and} \quad \gamma_{2,1} = 1 &\implies \frac{\partial}{\partial x}\mathrm{SS}(x) > 0 \quad \text{and} \quad \frac{\partial}{\partial x}\mathrm{mFPT}^{U_1}(x) < 0 \\
\gamma_{1,2} = 1 \quad \text{and} \quad \gamma_{2,1} > 1 &\implies \frac{\partial}{\partial x}\mathrm{SS}(x) < 0 \quad \text{and} \quad \frac{\partial}{\partial x}\mathrm{mFPT}^{U_1}(x) > 0 \\
\gamma_{1,2} < 1 \quad \text{and} \quad \gamma_{2,1} = 1 &\implies \frac{\partial}{\partial x}\mathrm{SS}(x) < 0 \quad \text{and} \quad \frac{\partial}{\partial x}\mathrm{mFPT}^{U_1}(x) > 0 \\
\gamma_{1,2} = 1 \quad \text{and} \quad \gamma_{2,1} < 1 &\implies \frac{\partial}{\partial x}\mathrm{SS}(x) > 0 \quad \text{and} \quad \frac{\partial}{\partial x}\mathrm{mFPT}^{U_1}(x) < 0,
\end{aligned}$$

for all $x > 0$. To prove this, we must consider the spanning trees and forests of $\mathcal{D}_N$, which can be used to generate analytical expressions for $\mathrm{SS}(x)$ (Eqns. 3 and 6) and $\mathrm{mFPT}^{U_1}(x)$ (Eqn. 7), as described in the main text and in Supporting Information S1. In particular, Eqn. 6 tells us that the vector, $\boldsymbol{\rho}(G)$, whose corresponding unit vector is the vector of steady-state probabilities, $\mathbf{p}^*(G)$, can be obtained by summing the products of edge labels of all spanning trees rooted at $j$, for each vertex $j \in \mathcal{V}(G)$. If we set $G = \mathcal{D}_N$, any such spanning tree can contain at most $N$ ligand-binding edges, labeled $k_{\mathrm{on}}x$, and therefore we have, for any $j \in \mathcal{V}(G)$,

$$\rho_j(\mathcal{D}_N) = a_0 + a_1 x + \cdots + a_N x^N,$$

where $a_0, \ldots, a_N$ are coefficients that do not depend on $x$. Subsequently normalizing by the coordinate sum of $\boldsymbol{\rho}(\mathcal{D}_N)$ (Eqn. 6) and applying Eqn. 3, it is easy to see that $\mathrm{SS}(x)$ must be a rational function of the form,

$$\mathrm{SS}(x) = \frac{a_{0,\mathrm{SS}} + a_{1,\mathrm{SS}}\, x + \cdots + a_{N,\mathrm{SS}}\, x^N}{b_{0,\mathrm{SS}} + b_{1,\mathrm{SS}}\, x + \cdots + b_{N,\mathrm{SS}}\, x^N}.$$

A similar argument can be made for $\mathrm{mFPT}^{U_1}(x)$, by appealing to Eqn. 7. Here, Eqn. 7 tells us that $\mathrm{mFPT}^{U_1}(x)$ can be calculated as a rational function in the edge labels of $\mathcal{D}_N^+$, where the numerator is the sum of edge label products of all spanning forests of $\mathcal{D}_N^+$ rooted at $\{j, M\}$, for each $j \in \mathcal{V}(\mathcal{D}_N)$, in which there is a path from $U_1$ to $j$; and the denominator is the sum of edge label products of all spanning trees of $\mathcal{D}_N^+$ rooted at $M$. Again, any such spanning forest and spanning tree must contain at most $N$ ligand-binding edges, and therefore we have,

$$\mathrm{mFPT}^{U_1}(x) = \frac{a_{0,\mathrm{mFPT}} + a_{1,\mathrm{mFPT}}\, x + \cdots + a_{N,\mathrm{mFPT}}\, x^N}{b_{0,\mathrm{mFPT}} + b_{1,\mathrm{mFPT}}\, x + \cdots + b_{N,\mathrm{mFPT}}\, x^N}.$$

Now, let $R(x)$ denote either of the two outputs, $\mathrm{SS}(x)$ or $\mathrm{mFPT}^{U_1}(x)$. Then differentiating $R$ with respect to $x$ yields,

$$\frac{\partial R}{\partial x} = \frac{\left(\sum_{i=1}^{N} a_{i,R}\, i x^{i-1}\right)\left(\sum_{i=0}^{N} b_{i,R}\, x^i\right) - \left(\sum_{i=0}^{N} a_{i,R}\, x^i\right)\left(\sum_{i=1}^{N} b_{i,R}\, i x^{i-1}\right)}{\left(\sum_{i=0}^{N} b_{i,R}\, x^i\right)^2}.$$

If we set $N = 2$, then this becomes

$$\frac{\partial R}{\partial x} = \frac{F(x)}{\left(b_{0,R} + b_{1,R}x + b_{2,R}x^2\right)^2},$$

where

$$F(x) = a_{1,R}b_{0,R} - a_{0,R}b_{1,R} + \left(2a_{2,R}b_{0,R} - 2a_{0,R}b_{2,R}\right)x + \left(a_{2,R}b_{1,R} - a_{1,R}b_{2,R}\right)x^2.$$

Since the denominator and $x$ are both positive, the sign of this derivative is determined by the signs of the coefficients of $F(x)$. For $N = 2$, these coefficients are easy to calculate explicitly. Assuming for now that $\gamma_{2,1} = 1$, we can write the coefficients of $R(x) = \mathrm{SS}(x)$ as,

$$\begin{aligned}
a_{1,\mathrm{SS}} b_{0,\mathrm{SS}} - a_{0,\mathrm{SS}} b_{1,\mathrm{SS}} &= k_{\mathrm{off}} k_{\mathrm{on}} \ell_{1,2} \ell_{2,1}^3 \left(\gamma_{1,2} - 1\right) + k_{\mathrm{off}}^3 k_{\mathrm{on}} \ell_{1,2} \ell_{2,1} \left(\gamma_{1,2} - 1\right) + k_{\mathrm{off}} k_{\mathrm{on}} \ell_{1,2}^3 \ell_{2,1} \left(\gamma_{1,2}^2 - \gamma_{1,2}\right) \\
&\quad + k_{\mathrm{off}} k_{\mathrm{on}} \ell_{1,2}^2 \ell_{2,1}^2 \left(\gamma_{1,2}^2 - 1\right) + k_{\mathrm{off}}^2 k_{\mathrm{on}} \ell_{1,2}^2 \ell_{2,1} \left(\gamma_{1,2}^2 - 1\right) + 2 k_{\mathrm{off}}^2 k_{\mathrm{on}} \ell_{1,2} \ell_{2,1}^2 \left(\gamma_{1,2} - 1\right) \\
a_{2,\mathrm{SS}} b_{0,\mathrm{SS}} - a_{0,\mathrm{SS}} b_{2,\mathrm{SS}} &= k_{\mathrm{off}} k_{\mathrm{on}}^2 \ell_{1,2} \ell_{2,1}^2 \left(\gamma_{1,2} - 1\right) + k_{\mathrm{off}}^2 k_{\mathrm{on}}^2 \ell_{1,2} \ell_{2,1} \left(\gamma_{1,2} - 1\right) + k_{\mathrm{off}} k_{\mathrm{on}}^2 \ell_{1,2}^2 \ell_{2,1} \left(\gamma_{1,2}^2 - \gamma_{1,2}\right) \\
a_{2,\mathrm{SS}} b_{1,\mathrm{SS}} - a_{1,\mathrm{SS}} b_{2,\mathrm{SS}} &= k_{\mathrm{off}} k_{\mathrm{on}}^3 \ell_{1,2} \ell_{2,1} \left(\gamma_{1,2} - 1\right).
\end{aligned}$$

Analogously, for $R(x) = \mathrm{mFPT}^{U_1}(x)$, we have

$$\begin{aligned}
a_{1,\mathrm{mFPT}} b_{0,\mathrm{mFPT}} - a_{0,\mathrm{mFPT}} b_{1,\mathrm{mFPT}} &= \ell_{1,2} k_{\mathrm{off}} k_{\mathrm{on}} r^4 \left(1 - \gamma_{1,2}\right) + \ell_{1,2} k_{\mathrm{off}}^3 k_{\mathrm{on}} r^2 \left(1 - \gamma_{1,2}\right) + \ell_{1,2}^2 k_{\mathrm{on}} r^4 \left(\gamma_{1,2} - \gamma_{1,2}^2\right) \\
&\quad + 2 \ell_{1,2} k_{\mathrm{off}}^2 k_{\mathrm{on}} r^3 \left(1 - \gamma_{1,2}\right) + \ell_{1,2} \ell_{2,1} k_{\mathrm{off}}^3 k_{\mathrm{on}} r \left(1 - \gamma_{1,2}\right) + \ell_{1,2} \ell_{2,1}^3 k_{\mathrm{off}} k_{\mathrm{on}} r \left(1 - \gamma_{1,2}\right) \\
&\quad + \ell_{1,2}^3 \ell_{2,1} k_{\mathrm{on}} r^2 \left(\gamma_{1,2} - \gamma_{1,2}^2\right) + 3 \ell_{1,2}^2 \ell_{2,1} k_{\mathrm{off}} k_{\mathrm{on}} r^2 \left(1 - \gamma_{1,2}^2\right) \\
&\quad + \ell_{1,2}^2 \ell_{2,1} k_{\mathrm{off}}^2 k_{\mathrm{on}} r \left(1 - \gamma_{1,2}^2\right) + \ell_{1,2}^2 \ell_{2,1}^2 k_{\mathrm{on}} r^2 \left(\gamma_{1,2} - \gamma_{1,2}^2\right) \\
&\quad + \ell_{1,2}^2 k_{\mathrm{off}}^2 k_{\mathrm{on}} r^2 \left(\gamma_{1,2} - \gamma_{1,2}^2\right) + \ell_{1,2}^2 \ell_{2,1}^2 k_{\mathrm{off}} k_{\mathrm{on}} r \left(1 - \gamma_{1,2}^2\right) \\
&\quad + 2 \ell_{1,2} \ell_{2,1}^2 k_{\mathrm{off}}^2 k_{\mathrm{on}} r \left(1 - \gamma_{1,2}\right) + 2 \ell_{1,2}^2 \ell_{2,1} k_{\mathrm{on}} r^3 \left(\gamma_{1,2} - \gamma_{1,2}^2\right) \\
&\quad + 2 \ell_{1,2}^2 k_{\mathrm{off}} k_{\mathrm{on}} r^3 \left(\gamma_{1,2} - \gamma_{1,2}^2\right) + 3 \ell_{1,2} \ell_{2,1} k_{\mathrm{off}} k_{\mathrm{on}} r^3 \left(1 - \gamma_{1,2}\right) \\
&\quad + 3 \ell_{1,2} \ell_{2,1}^2 k_{\mathrm{off}} k_{\mathrm{on}} r^2 \left(1 - \gamma_{1,2}\right) + 4 \ell_{1,2} \ell_{2,1} k_{\mathrm{off}}^2 k_{\mathrm{on}} r^2 \left(1 - \gamma_{1,2}\right) \\
&\quad + \ell_{1,2}^3 \ell_{2,1} k_{\mathrm{off}} k_{\mathrm{on}} r \left(\gamma_{1,2} - \gamma_{1,2}^2\right) \\
a_{2,\mathrm{mFPT}} b_{0,\mathrm{mFPT}} - a_{0,\mathrm{mFPT}} b_{2,\mathrm{mFPT}} &= 2 \ell_{1,2} k_{\mathrm{off}} k_{\mathrm{on}}^2 r^3 \left(1 - \gamma_{1,2}\right) + 2 \ell_{1,2} k_{\mathrm{off}}^2 k_{\mathrm{on}}^2 r^2 \left(1 - \gamma_{1,2}\right) + 2 \ell_{1,2}^2 k_{\mathrm{on}}^2 r^3 \left(\gamma_{1,2} - \gamma_{1,2}^2\right) \\
&\quad + 2 \ell_{1,2} \ell_{2,1} k_{\mathrm{off}}^2 k_{\mathrm{on}}^2 r \left(1 - \gamma_{1,2}\right) + 2 \ell_{1,2} \ell_{2,1}^2 k_{\mathrm{off}} k_{\mathrm{on}}^2 r \left(1 - \gamma_{1,2}\right) \\
&\quad + 2 \ell_{1,2}^2 \ell_{2,1} k_{\mathrm{on}}^2 r^2 \left(\gamma_{1,2} - \gamma_{1,2}^2\right) + 2 \ell_{1,2}^2 k_{\mathrm{off}} k_{\mathrm{on}}^2 r^2 \left(\gamma_{1,2} - \gamma_{1,2}^2\right) \\
&\quad + 4 \ell_{1,2} \ell_{2,1} k_{\mathrm{off}} k_{\mathrm{on}}^2 r^2 \left(1 - \gamma_{1,2}\right) + 2 \ell_{1,2}^2 \ell_{2,1} k_{\mathrm{off}} k_{\mathrm{on}}^2 r \left(\gamma_{1,2} - \gamma_{1,2}^2\right) \\
a_{2,\mathrm{mFPT}} b_{1,\mathrm{mFPT}} - a_{1,\mathrm{mFPT}} b_{2,\mathrm{mFPT}} &= \ell_{1,2} k_{\mathrm{off}} k_{\mathrm{on}}^3 r^2 \left(1 - \gamma_{1,2}\right) + \ell_{1,2}^2 k_{\mathrm{on}}^3 r^2 \left(\gamma_{1,2} - \gamma_{1,2}^2\right) + \ell_{1,2} \ell_{2,1} k_{\mathrm{off}} k_{\mathrm{on}}^3 r \left(1 - \gamma_{1,2}\right).
\end{aligned}$$

Since all edge labels are positive, these expressions clearly show that signs of $(\partial/\partial x)\,\mathrm{SS}$ and $(\partial/\partial x)\,\mathrm{mFPT}^{U_1}$ are determined by the value of $\gamma_{1,2}$. Namely, if $\gamma_{1,2} > 1$, then all three coefficients of $(\partial/\partial x)\,\mathrm{SS}$ must be positive, whereas the coefficients of $(\partial/\partial x)\,\mathrm{mFPT}^{U_1}$ must be negative. Similarly, if $\gamma_{1,2} < 1$, then all three coefficients of $(\partial/\partial x)\,\mathrm{SS}$ are negative, whereas the coefficients of $(\partial/\partial x)\,\mathrm{mFPT}^{U_1}$ are positive. This proves the desired claim for $N = 2$. An analogous argument can be made for $\gamma_{2,1}$.

For $N = 3$, a similar set of arguments can be used to show that, when the ligand regulates at most two of the ligand-bound transitions ($B_1 \to B_2$, $B_2 \to B_1$, $B_2 \to B_3$, and $B_3 \to B_2$), then the signs of $(\partial/\partial x)\,\mathrm{SS}(x)$ and $(\partial/\partial x)\,\mathrm{mFPT}^{U_1}(x)$ follow the signs given in Table 1. (Here, the "or" in each condition should be interpreted as inclusive, e.g., the first condition requires that either $\gamma_{1,2} > 1$ and $\gamma_{2,3} = 1$, or $\gamma_{1,2} = 1$ and $\gamma_{2,3} > 1$, or $\gamma_{1,2}, \gamma_{2,3} > 1$.) We used Mathematica to calculate the formulas for the coefficients in $\mathrm{SS}(x)$ and $\mathrm{mFPT}^{U_1}(x)$, which are much more complicated than for the $N = 2$ case; given this complexity, we do not provide them here, but they can be found in ċsv files in the GitHub repository: `https://github.com/theobiolab/FPT_paper.git`.

### S3 Impossibility of decoupling with coherent regulation and equal transition rates in $\mathcal{D}_3$

Here, we show that decoupling is not achievable in the ladder model, $\mathcal{D}_3$, with any coherent regulatory regime in which

$$\gamma_{1,2} \geq 1, \qquad \gamma_{2,3} \geq 1, \qquad \gamma_{2,1} = \gamma_{3,2} = 1, \tag{S10}$$

when we are subject to the constraint,

$$\ell_{1,2} = \ell_{2,1} = \ell_{2,3} = \ell_{3,2}. \tag{S11}$$

| Condition | Sign of $(\partial/\partial x)\,\mathrm{SS}(x)$ | Sign of $(\partial/\partial x)\,\mathrm{mFPT}^{U_1}(x)$ |
|---|---|---|
| $\gamma_{1,2} > 1$ or $\gamma_{2,3} > 1$, $\gamma_{2,1} = \gamma_{3,2} = 1$ | $+$ | $-$ |
| $\gamma_{2,1} > 1$ or $\gamma_{3,2} > 1$, $\gamma_{1,2} = \gamma_{2,3} = 1$ | $-$ | $+$ |
| $\gamma_{1,2} < 1$ or $\gamma_{2,3} < 1$, $\gamma_{2,1} = \gamma_{3,2} = 1$ | $-$ | $+$ |
| $\gamma_{2,1} < 1$ or $\gamma_{3,2} < 1$, $\gamma_{1,2} = \gamma_{2,3} = 1$ | $+$ | $-$ |
| $\gamma_{1,2} > 1$ or $\gamma_{3,2} < 1$, $\gamma_{2,1} = \gamma_{2,3} = 1$ | $+$ | $-$ |
| $\gamma_{1,2} < 1$ or $\gamma_{3,2} > 1$, $\gamma_{2,1} = \gamma_{2,3} = 1$ | $-$ | $+$ |
| $\gamma_{2,1} < 1$ or $\gamma_{2,3} > 1$, $\gamma_{1,2} = \gamma_{3,2} = 1$ | $+$ | $-$ |
| $\gamma_{2,1} > 1$ or $\gamma_{2,3} < 1$, $\gamma_{1,2} = \gamma_{3,2} = 1$ | $-$ | $+$ |

**Table 1.** Signs of $(\partial/\partial x)\,\mathrm{SS}(x)$ and $(\partial/\partial x)\,\mathrm{mFPT}^{U_1}(x)$ in the ladder model $\mathcal{D}_3$, for each of the specified regulatory regimes, for all $x > 0$.

In particular, we shall show that, in this regime,

$$\Delta_{\overline{\mathrm{SS}}} < \Delta_{\overline{\mathrm{mFPT}}^{U_1}}. \tag{S12}$$

Using symbolic calculations to evaluate the two dynamic ranges with $\ell_{1,2} = \ell_{2,1}$ and $\ell_{2,3} = \ell_{3,2}$ (Table 2, case 2.I), taking the difference, and additionally setting $\alpha = \ell_{2,3}/\ell_{1,2} = 1$, we get

$$\Delta_{\overline{\mathrm{SS}}} - \Delta_{\overline{\mathrm{mFPT}}^{U_1}} = \frac{2\gamma_{1,2}\gamma_{2,3} - \gamma_{1,2} - 1}{3\left(\gamma_{1,2}\gamma_{2,3} + \gamma_{1,2} + 1\right)} - \frac{3\gamma_{1,2}\gamma_{2,3}\beta + 2\gamma_{1,2}\gamma_{2,3} - \gamma_{1,2}\beta - \gamma_{1,2} - \gamma_{2,3}\beta - \beta - 1}{3\gamma_{1,2}\gamma_{2,3}\left(\beta + 1\right)} = \frac{P}{Q},$$

where

$$\begin{aligned} P = &-\gamma_{1,2}^2\gamma_{2,3}^2\beta - 3\gamma_{1,2}^2\gamma_{2,3}\beta - 2\gamma_{1,2}^2\gamma_{2,3} + \gamma_{1,2}^2\beta + \gamma_{1,2}^2 + \gamma_{1,2}\gamma_{2,3}^2\beta - 2\gamma_{1,2}\gamma_{2,3}\beta - 2\gamma_{1,2}\gamma_{2,3} + 2\gamma_{1,2}\beta \\ &+ 2\gamma_{1,2} + \gamma_{2,3}\beta + \beta + 1 \\ Q = &\,3\left(\gamma_{1,2}^2\gamma_{2,3}^2\beta + \gamma_{1,2}^2\gamma_{2,3}^2 + \gamma_{1,2}^2\gamma_{2,3}\beta + \gamma_{1,2}^2\gamma_{2,3} + \gamma_{1,2}\gamma_{2,3}\beta + \gamma_{1,2}\gamma_{2,3}\right). \end{aligned}$$

We want to show that this difference is negative whenever $\gamma_{1,2}, \gamma_{2,3} \geq 1$. Since the denominator, $Q$, is positive, we focus on the numerator, $P$. If we define $\varepsilon = \gamma_{2,3} - 1 \geq 0$, we can rewrite $P$ as

$$P = -\gamma_{1,2}^2\beta\varepsilon^2 - 5\gamma_{1,2}^2\beta\varepsilon - 3\gamma_{1,2}^2\beta - 2\gamma_{1,2}^2\varepsilon - \gamma_{1,2}^2 + \gamma_{1,2}\beta\varepsilon^2 + \gamma_{1,2}\beta - 2\gamma_{1,2}\varepsilon + \beta\varepsilon + 2\beta + 1, \tag{S13}$$

which we can rearrange further as

$$P = \gamma_{1,2}\beta\varepsilon^2\left(1 - \gamma_{1,2}\right) + \beta\varepsilon\left(1 - 5\gamma_{1,2}^2\right) + \beta\left(\gamma_{1,2} + 2 - 3\gamma_{1,2}^2\right) + \left(1 - \gamma_{1,2}^2\right) - 2\varepsilon\gamma_{1,2}\left(1 + \gamma_{1,2}\right).$$

From here, it is easy to see that, since $\gamma_{1,2} \geq 1$, the first, third, and fourth terms are less than or equal to zero, and the second and fifth terms must be less than zero. In particular, we have

$$\begin{aligned} P &\leq \beta\varepsilon\left(1 - 5\gamma_{1,2}^2\right) - 2\varepsilon\gamma_{1,2}\left(1 + \gamma_{1,2}\right) \\ &\leq -4\beta\varepsilon - 4\varepsilon \\ &< 0, \end{aligned}$$

from which Eqn. S12 follows.

### S4 Decoupling via regulation of backward transitions in $\mathcal{D}_3$

Here, we describe the optimisation procedure for regulatory regimes in which the ligand regulates one or both of the backward transitions, $B_2 \to B_1$ and $B_3 \to B_2$, in the ladder model, $\mathcal{D}_3$. The corresponding results are given in Figs. S6, S7, and S8.

We assumed that the ligand regulates one of each pair of reversible transitions, i.e., that it regulates either $B_1 \to B_2$ or $B_2 \to B_1$, and that it regulates either $B_2 \to B_3$ or $B_3 \to B_2$. As such, we considered three regulatory regimes:

1. $\gamma_{2,1}, \gamma_{2,3} \neq 1$ and $\gamma_{1,2} = \gamma_{3,2} = 1$ (Fig. S6);
2. $\gamma_{1,2}, \gamma_{3,2} \neq 1$ and $\gamma_{2,1} = \gamma_{2,3} = 1$ (Fig. S7); and
3. $\gamma_{2,1}, \gamma_{3,2} \neq 1$ and $\gamma_{1,2} = \gamma_{2,3} = 1$ (Fig. S8),

to complement the analyses of the case in which $B_1 \to B_2$ and $B_2 \to B_3$ are regulated $(\gamma_{1,2}, \gamma_{2,3} \neq 1, \gamma_{2,1} = \gamma_{3,2} = 1)$ on which we have focused for the majority of this paper. To simplify the analysis, we additionally set (Figs. S6A, S7A, and S8A)

$$\ell_f = \ell_{1,2} = \ell_{2,3} = r \qquad \text{and} \qquad \ell_b = \ell_{2,1} = \ell_{3,2}.$$

We then used PSO as previously described to solve the following optimisation problem:

$$\begin{aligned} \text{minimize} \quad & f = 1 - \left(\Delta_{\overline{\text{SS}}} - \Delta_{\overline{\text{mFPT}^{U_1}}}\right) \\ \text{subject to} \quad & g_1 = \left|\log_{10} \gamma_{i,j}\right| - k \leq 0 \\ & g_2 = \left|\log_{10} \gamma_{i',j'}\right| - k \leq 0, \end{aligned}$$

where $\gamma_{i,j}$ and $\gamma_{i',j'}$ are the two nontrivial regulatory factors in each regime, and the rates $\ell_f$, $\ell_b$, $k_{\text{off}}$, and $k_{\text{on}}$ were restricted to lie in the range $\left[10^{-4}, 10^4\right]$ in the appropriate units (units of $\delta$ for $\ell_f$, $\ell_b$, and $k_{\text{off}}$, and units of $\delta / (1\,\text{c.u.})$ for $k_{\text{on}}$). The optimizations in Fig. S6 S7 S8 are performed setting $k = 3$.

### S5 Derivation of the activation time $\text{mFPT}^{\langle U \rangle}$ (Eqn. 26)

Here, we provide a derivation of the activation time, $\text{mFPT}^{\langle U \rangle}$, from an equilibrium of initial states (Eqn. 26). Using the notation and terminology established in the main text and in Supplemental Information S1, we define this activation time as the mFPT to produce one *additional* molecule of the readout, $M$, averaged over all possible initial system states (i.e., the graph vertices) and readout copy-numbers. If we denote by $X(t)$ and $n_M(t)$ the system state and readout copy-number at time $t$, respectively, we may write this quantity as

$$\text{mFPT}^{\langle U \rangle}(x) = \sum_{n=0}^{\infty} \sum_{i \in \mathcal{V}(G)} p^*_{i,n}(x = 0) \cdot \mathbb{E}\left[\inf\{t > 0 : n_M(t) = n + 1\} \mid X(0) = i \text{ and } n_M(0) = n\right]. \tag{S14}$$

Now, in defining the infinite "copy-number graph" that describes this composite Markov process (Supplemental Information S1) [6], we impose the assumption that the transition rates between two system states, $i, j \in \mathcal{V}(G)$, does not depend on the readout copy-number. This implies that the mFPT in the above right-hand sum does not depend on $n_M(0) = n$, and is in fact merely given by

$$\mathbb{E}\left[\inf\{t > 0 : n_M(t) = n + 1\} \mid X(0) = i \text{ and } n_M(0) = n\right] = \text{mFPT}^i(x).$$

Therefore, we can rewrite Eqn. S14 as

$$\text{mFPT}^{\langle U \rangle} = \sum_{i \in \mathcal{V}(G)} \text{mFPT}^i(x) \sum_{n=0}^{\infty} p^*_{i,n}(x = 0).$$

Now, we recall from Supplemental Information S1 that

$$\sum_{n=0}^{\infty} p^*_{i,n} = q^*_i = p^*_i,$$

so that we may simply write

$$\text{mFPT}^{\langle U \rangle} = \sum_{i \in \mathcal{V}(G)} \left(p^*_i(x = 0) \cdot \text{mFPT}^i(x)\right).$$

Now, setting $G = \mathcal{D}_N$, we note that the probability of every ligand-bound vertex is zero in the absence of ligand, i.e., $p^*_{B_i}(x=0) = 0$ for all $i = 1, \ldots, N$. In this case, we get

$$\mathrm{mFPT}^{\langle U \rangle} = \sum_{i=1}^{N} \left( p^*_{U_i}(x=0) \cdot \mathrm{mFPT}^{U_i}(x) \right),$$

i.e., we recover Eqn. 26.

### S6 A simple model for *hunchback* regulation by Bicoid and Zelda, related to [11]

In [11], Eck *et al.* examined the effects of the TFs Bicoid (Bcd) and Zelda (Zld) on the transcription of a reporter driven by the well-known *hunchback* minimal enhancer P2 in the *Drosophila* blastoderm. In this system, Bcd exhibits a concentration gradient over the antero-posterior axis of the embryo, whereas Zld concentration is essentially constant [11]. The authors examined the activation time and levels of the reporter. In WT embryos, the activation time was constant throughout the embryo while the transcription levels varied (thus exhibiting decoupling). In Zld null embryos both quantities became coupled. Here, we provide a highly simplified model for this system, in which Zld is modelled implicitly and Bcd is described to simply bind to one regulatory site. The goal is to show how our formalism can be easily adapted to account for specific experimental systems and data, although the construction of a detailed model, with multiple TFs and binding sites and appropriate parameterization, is outside the scope of the present paper.

We use a variation of the ladder model $\mathcal{D}_3$, as shown in Fig. S11, in which Bcd serves as the ligand, and the system transits through three internal regulatory states before producing the molecular readout, which is the reporter mRNA. The first set of transitions, $U_1 \rightleftharpoons U_2$ and $B_1 \rightleftharpoons B_2$, represent chromatin dynamics, whereas the second set of transitions, $U_2 \rightleftharpoons U_3$ and $B_2 \rightleftharpoons B_3$, represent RNA polymerase recruitment and initiation. For simplicity, we assume that each of these sets of transitions is governed by a characteristic timescale, namely the rates $\ell_1$ for the first set and $\ell_2$ for the second (Fig. S11), and assume that $\ell_2 \gg \ell_1$. To model Zld's role as a pioneer factor that opens inaccessible chromatin and enables other TFs, including Bcd, to access DNA [11,12], we assume that it contributes to regulating both $U_1 \to U_2$ and $B_1 \to B_2$, via the regulatory factor $\gamma_{zld} \geq 1$. To model Bcd's role as a TF that activates *hunchback* transcription, we assume that it contributes to regulating both $B_1 \to B_2$ and $B_2 \to B_3$, via the regulatory factor $\gamma_{bcd} > 1$. Finally, we assume that the regulatory effects of Zld and Bcd on $B_1 \to B_2$ are additive, so that its transition rate is given by $(\gamma_{zld} + \gamma_{bcd})\,\ell_1$.

We first consider the case where Zld is absent, so that $\gamma_{zld} = 1$. In this case, the model reduces to an instance of case 2.I (Table 2), but with $\gamma_{1,2} = \gamma_{2,3} = \gamma_{bcd}$. As implied by our results in Fig. 2F–J, in this regime, the steady-state level and activation time of the gene are coupled. This can also be observed by directly appealing to the corresponding formula for $\Delta_{\overline{\mathrm{mFPT}^{U_1}}}$ in Table 2. Setting $\gamma_{1,2} = \gamma_{2,3} = \gamma_{bcd}$ and $\alpha = \beta = \ell_2/\ell_1$, we obtain,

$$\Delta_{\overline{\mathrm{mFPT}^{U_1}}} = -\frac{\alpha^2 \gamma_{bcd}\left(\gamma_{bcd} - 1\right) + 2\alpha\left(2\gamma_{bcd}^2 - \gamma_{bcd} - 1\right)}{\gamma_{bcd}^2\left(\alpha^2 + 5\alpha\right)}.$$

Since $\gamma_{bcd} > 1$, both terms in the numerator must be positive; therefore, $\Delta_{\overline{\mathrm{mFPT}^{U_1}}} < 0$. Indeed, this is what is found by Eck *et al.* in a *zelda*$^-$ mutant background.

We now turn to the case where Zld is present, so that $\gamma_{zld} \neq 1$. In this case, we obtain the following dynamic ranges for the steady-state and activation time for $\ell_2 \gg \ell_1$:

$$\Delta_{\overline{\mathrm{SS}}} = \frac{\gamma_{zld} + 1 + \dfrac{\gamma_{zld}^2}{\gamma_{bcd}} - \dfrac{\gamma_{zld}^2}{\gamma_{bcd}^2} - \dfrac{\gamma_{zld}}{\gamma_{bcd}^2}}{2\gamma_{zld} + 1 + \dfrac{2\gamma_{zld}^2}{\gamma_{bcd}} + \dfrac{3\gamma_{zld}}{\gamma_{bcd}} + \dfrac{1}{\gamma_{bcd}} + \dfrac{2\gamma_{zld}^2}{\gamma_{bcd}^2} + \dfrac{3\gamma_{zld}}{\gamma_{bcd}^2} + \dfrac{1}{\gamma_{bcd}^2}}$$

$$\Delta_{\overline{\mathrm{mFPT}^{U_1}}} = -\frac{1}{1 + \dfrac{\gamma_{zld}}{\gamma_{bcd}}}.$$

These equations show that, in the limit where the regulatory effect provided by Zld is much greater than that of Bcd ($\gamma_{zld} \gg \gamma_{bcd}$) and the first set of transitions is much slower than the second ($\ell_2 \gg \ell_1$), we obtain decoupling: $\Delta_{\overline{\mathrm{mFPT}}^{U_1}}$ goes to zero while $\Delta_{\overline{\mathrm{SS}}}$ tends to a finite, nonzero value, again in line with the observations of [11].

### S7 Notes on the Erlang process, related to [13]

In [13], Alamos *et al.* employed a simple "kinetic barrier" model to describe the onset of transcription after mitosis in early *Drosophila* development. In this model, a synthetic enhancer that harbors one binding site for the morphogen Dorsal (Dl) regulates transcription of a reporter gene, by irreversibly transitioning through a sequence of transcriptionally inactive states, $\mathrm{OFF}_1, \ldots, \mathrm{OFF}_n$, to a terminal, transcriptionally active state, ON, with each transition proceeding with the same rate, $k([\mathrm{Dl}])$, which is given by

$$k([\mathrm{Dl}]) = c\left(\frac{[\mathrm{Dl}]/K_d}{1 + [\mathrm{Dl}]/K_d}\right),$$

where $c$ is a basal rate and $K_d$ is the Dl–DNA dissociation constant. Once in the ON state, the steady-state transcription level is defined in terms of a separate "thermodynamic model" that incorporates Dl and RNA polymerase binding. The mFPT from $\mathrm{OFF}_1$ to ON was used as a measure of the activation time.

Interestingly, Alamos *et al.* used this model to explain decoupling between steady-state transcription level and activation time with respect to Dl concentration, but without rate scale separation among the transition rates. This appears to be at odds with our results, which suggest that a system in which a TF coherently promotes production of the readout can exhibit decoupling only when there is rate scale separation (Fig. 2). Here, we provide one possible explanation for this apparent discrepancy, as arising from the fact that Alamos *et al.* used a short time window to estimate the mFPT to the ON state. This contrasts with our definition of the mFPT, which is not subject to such truncation.

Alamos *et al.*'s model is an instance of the well-characterised Erlang process, which is a Markov process on a set of $N+1$ states, $0, \ldots, N$, with a linear sequence of irreversible transitions, $i \to i+1$ for $i = 0, \ldots, N-1$, each with the same transition rate, $k$. Alamos *et al.* defined the rate $k$ as being time-dependent, because the Dl concentration is dynamic over the observation window. However, for computations of input-output responses, Alamos *et al.* used Dl concentration measurements from a single timepoint, around 7 minutes into nuclear cycle 13 [13, Fig. S8B]. This model can be schematically depicted using the following graph:

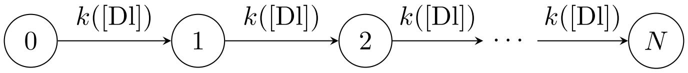


Alamos *et al.* defined the "mean transcription onset time," which corresponds to what we call the activation time, as

$$\mathbb{E}[\mathrm{onset}] = \sum_{i=1}^{m} t_i p_i, \tag{S15}$$

where $t_i$ is one of a set of $m$ possible onset times, $t_1, \ldots, t_m = T$, and $p_i$ is the probability of observing the onset time $t_i$. (Note that Alamos *et al.* modeled time as a discrete variable, which advances in units of a small timestep $dt$.) Crucially, Alamos *et al.* assumed that the onset time is capped at some maximal time $t_m = T = 7\,\mathrm{min}$, which represents the time window over which transcriptional activation at the reporter gene was observed in each nucleus. As such, the probabilities $p_i$ are normalized over this window, as

$$\sum_{i=1}^{m} p_i = 1. \tag{S16}$$

Now, the FPT from state 0 to state $N$ in this process is well-known to follow the probability distribution function,

$$f(t) = \frac{k^N t^{N-1} e^{-kt}}{(N-1)!},$$

where we have dropped the dependence of $k$ on [Dl] for clarity. The corresponding cumulative distribution function is given by

$$F(t') = \int_0^{t'} f(t)\,dt = 1 - \sum_{n=0}^{N-1} \frac{(kt')^n\, e^{-kt'}}{n!};$$

this function describes the fraction of nuclei in which transcription has been activated by time $t$. Note that, since $f(t)$ is a probability distribution on $[0, \infty)$,

$$F(T) = \int_0^T f(t)\,dt < 1.$$

To follow Alamos *et al.*'s assumption and truncate the FPT distribution onto the domain $[0, T]$, we must normalise $f(t)$ by $F(T)$, to get

$$f^{\text{trunc}}(t; T) = \frac{f(t)}{F(T)}.$$

From here, we can define the onset time, as per Alamos *et al.*, as the integral,

$$\mathbb{E}[\text{onset}] = \int_0^T t f^{\text{trunc}}(t; T)\,dt = \frac{1}{F(T)} \int_0^T t \left( \frac{k^N t^{N-1} e^{-kt}}{(N-1)!} \right) dt.$$

Integrating by parts, this can be evaluated as

$$\begin{aligned}
\mathbb{E}[\text{onset}] &= \frac{k^N}{F(T)\,(N-1)!} \int_0^T t^N e^{-kt} dt \\
&= \frac{k^N}{F(T)\,(N-1)!} \left( \frac{N!}{k^{N+1}} - \sum_{n=0}^{N} \left( \frac{N!}{k^{n+1}\,(N-n)!} \right) T^{N-n} e^{-kT} \right) \\
&= \frac{k^N}{F(T)\,(N-1)!} \left( \frac{N!}{k^{N+1}} \right) \left( 1 - \sum_{n=0}^{N} \frac{k^{N-n} T^{N-n} e^{-kT}}{(N-n)!} \right) \\
&= \frac{N}{kF(T)} \left( 1 - \sum_{n=0}^{N} \frac{(kT)^n\, e^{-kT}}{n!} \right).
\end{aligned} \tag{S17}$$

This is distinct from the mean of the untruncated FPT distribution, which is given by

$$\int_0^\infty t f(t)\,dt = \frac{k^N}{(N-1)!} \int_0^\infty t^N e^{-kt}\,dt = \frac{N}{k}. \tag{S18}$$

Now, we sought to compare the untruncated and truncated FPT distributions, $f(t)$ and $f^{\text{trunc}}(t; T)$, and their means, given in Eqns. S18 and S17, respectively (Fig. S12A). To follow Alamos *et al.*'s analysis as closely as possible, we assumed that $T = 7\,\text{min}$, and set the coefficients in $k([\text{Dl}])$ to $c = 0.55\,\text{min}^{-1}$ and $K_d = 250\,\text{a.u.}$ [13]. We observed that the untruncated FPT distribution exhibits a strong dependence on the Dl concentration for various choices of $N$ (Fig. S12A). In contrast, the truncated FPT distribution shows a much weaker dependence on the Dl concentration, especially for $N = 6$ (Fig. S12B–C). This distinction is also visible between the means, as shown in Fig. S12D: the untruncated mean decays sharply with Dl concentration for various values of $N$, whereas the truncated mean—which corresponds to Alamos *et al.*'s definition of onset time—is largely independent of the Dl concentration, especially for $N = 6$. As we describe in the Discussion, these results suggest that Alamos *et al.*'s choice of model only captures the observed decoupling if it enforces a finite time window during which transcription can occur.

### S8 Definition of activation time according to the master equation

As described in Supplemental Information S1, the copy-number $n_M$ of the molecular readout $M$ can be viewed as a random variable that is governed by its own master equation (Eqn. S3). In some other works in the literature, the activation time is defined as the time required for the mean copy-number, $\langle n_M \rangle$, to reach a given threshold value [14,15]. Here, we describe how this definition differs from the mFPT that we have used throughout this paper.

Let us consider the random telegraph model, $\mathcal{C}_2$, in Fig. 1E, with $\ell_{1,2}(x) = a(x)$ and $\ell_{2,1}(x) = b(x)$ for ease of notation. As described in the main text, the mFPT from 1 to $M$ in the augmented graph, $\mathcal{C}_2^+$, which measures the mFPT to the production of one molecule of $M$, is given by (Eqn. 15)

$$\text{mFPT}^1(x) = \frac{a+b+r}{ar}. \tag{S19}$$

Meanwhile, the master equation for the mean copy-number, $\langle n_M \rangle$, is given by

$$\frac{d}{dt}\langle n_M \rangle = \frac{d}{dt}\sum_{i=1}^{2} \mu_i(t),$$

where $\boldsymbol{\mu}(t) = (\mu_1(t), \mu_2(t))^{\mathrm{T}}$ is the vector defined in Supplemental Information S1. Using Eqn. S3, we can write a master equation for this vector, as

$$\begin{aligned}
\frac{d}{dt}\begin{bmatrix} \mu_1(t) \\ \mu_2(t) \end{bmatrix} &= \underbrace{\begin{bmatrix} 0 & 0 \\ 0 & r \end{bmatrix}}_{=\mathbf{R}} \underbrace{\begin{bmatrix} q_1(t) \\ q_2(t) \end{bmatrix}}_{=\mathbf{q}(t)} - \delta \begin{bmatrix} \mu_1(t) \\ \mu_2(t) \end{bmatrix} + \underbrace{\begin{bmatrix} -a & b \\ a & -b \end{bmatrix}}_{=\mathcal{L}(\mathcal{C}_2)} \begin{bmatrix} \mu_1(t) \\ \mu_2(t) \end{bmatrix} \\
&= \begin{bmatrix} 0 \\ rq_2(t) \end{bmatrix} + \begin{bmatrix} -\delta - a & b \\ a & -\delta - b \end{bmatrix} \begin{bmatrix} \mu_1(t) \\ \mu_2(t) \end{bmatrix},
\end{aligned}$$

where $q_1(t)$ and $q_2(t)$ are the time-dependent probabilities of vertices 1 and 2 in $\mathcal{C}_2$, respectively. From here, we get

$$\begin{aligned}
\frac{d\langle n_M \rangle}{dt} &= \frac{d\mu_1}{dt} + \frac{d\mu_2}{dt} \\
&= rq_2(t) - \delta\left(\mu_1(t) + \mu_2(t)\right) \\
&= rq_2(t) - \delta\langle n_M \rangle.
\end{aligned} \tag{S20}$$

In turn, $q_2(t)$ satisfies the master equation,

$$\frac{d}{dt}\begin{bmatrix} q_1(t) \\ q_2(t) \end{bmatrix} = \underbrace{\begin{bmatrix} -a & b \\ a & -b \end{bmatrix}}_{=\mathcal{L}(\mathcal{C}_2)} \begin{bmatrix} q_1(t) \\ q_2(t) \end{bmatrix}. \tag{S21}$$

Now, assuming initial values of $q_1(0) = 1$ and $q_2(0) = 0$, it is straightforward to solve for the time-dependent solution for Eqn. S21 from the eigenvalues and eigenvectors of $\mathcal{L}(\mathcal{C}_2)$, as

$$\begin{bmatrix} q_1(t) \\ q_2(t) \end{bmatrix} = \left(\frac{1}{a+b}\right)\begin{bmatrix} b \\ a \end{bmatrix} + \left(\frac{1}{a+b}\right) e^{-(a+b)t} \begin{bmatrix} a \\ -a \end{bmatrix},$$

which we can substitute into Eqn. S20 to get

$$\frac{d\langle n_M \rangle}{dt} = \left(\frac{ra}{a+b}\right)\left(1 - e^{-(a+b)t}\right) - \delta\langle n_M \rangle.$$

This can be rearranged as

$$\frac{d\langle n_M \rangle}{dt} + \delta\langle n_M \rangle = \left(\frac{ra}{a+b}\right)\left(1 - e^{-(a+b)t}\right),$$

which is a first-order differential equation that can be solved by way of an integrating factor of $e^{\delta t}$. This yields the general solution,

$$\begin{aligned}\langle n_M \rangle &= e^{-\delta t}\left(\int e^{\delta t}\left(\frac{ra}{a+b}\right)\left(1-e^{-(a+b)t}\right)dt + C\right)\\ &= e^{-\delta t}\left(\left(\frac{ra}{a+b}\right)\left(\frac{e^{\delta t}}{\delta} - \frac{e^{-(a+b-\delta)t}}{\delta-a-b}\right) + C\right)\\ &= \left(\frac{ra}{a+b}\right)\left(\frac{1}{\delta} - \frac{e^{-(a+b)t}}{\delta-a-b}\right) + Ce^{-\delta t}.\end{aligned}$$

Assuming an initial value of $\langle n_M \rangle(0) = 0$, we obtain the specific solution,

$$\langle n_M \rangle = \left(\frac{ra}{a+b}\right)\left(\frac{1}{\delta} - \frac{e^{-(a+b)t}}{\delta-a-b}\right) - \left(\frac{ra}{a+b}\right)\left(\frac{1}{\delta} - \frac{1}{\delta-a-b}\right)e^{-\delta t}. \tag{S22}$$

We can then define the activation time as the time by which $\langle n_M \rangle$ first reaches a threshold value, say, $\langle n_M \rangle = 1$.

This solution makes clear what was already suggested in Eqn. S20: this definition of activation time depends on the degradation rate, $\delta$ (Fig. S10A). Meanwhile, the mFPT in Eqn. S19 does not depend on $\delta$ (Fig. S10B). This is because the two measures treat degradation in fundamentally different ways. The former measure, based on $\langle n_M \rangle$, simply quantifies the time to which the copy-number of $M$ reaches 1 on average. Since $M$ is dynamically produced and degraded over time, this measure naturally depends on $\delta$. The latter measure, based on $\mathrm{mFPT}^1(x)$, quantifies the time to which the *regulatory system* produces one copy of $M$, regardless of how many copies of $M$ are present in the environment. As such, this measure merely depends on the internal dynamics of the regulatory system and its ligand-binding state, while ignoring the degradation of $M$ as an entirely separate process that takes place elsewhere. As how these internal dynamics can give rise to decoupling constitutes our main topic of interest in this paper, we chose to focus on the mFPT-based measure; however, we believe that both measures represent reasonable interpretations of activation time (Discussion).

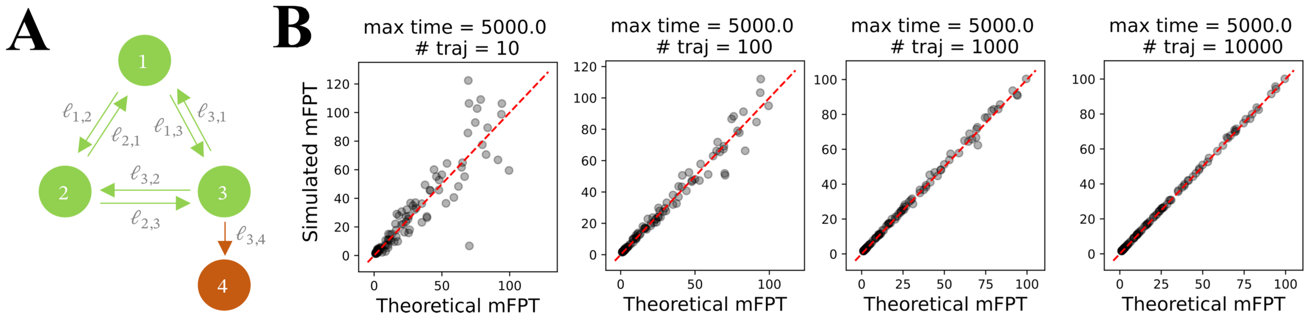


**Fig S1. Agreement between mFPT from the Chebotarev–Agaev recurrence (Eqn. 12) and estimates from Gillespie simulations**. **(A)** Three-state model with one absorbing state. Each edge label represents an infinitesimal transition rate of the embedded Markov process. **(B)** Agreement between the theoretically derived mFPT and estimates using a Python implementation of the Gillespie algorithm, GillesPy2 [16]. Parameter values were sampled from a log-uniform distribution on $[10^{-3}, 10^{3}]$. "Max time" refers to the maximum simulation time, "# traj" to the number of simulated trajectories. The red dashed line represents the line where theoretical and simulated values coincide.

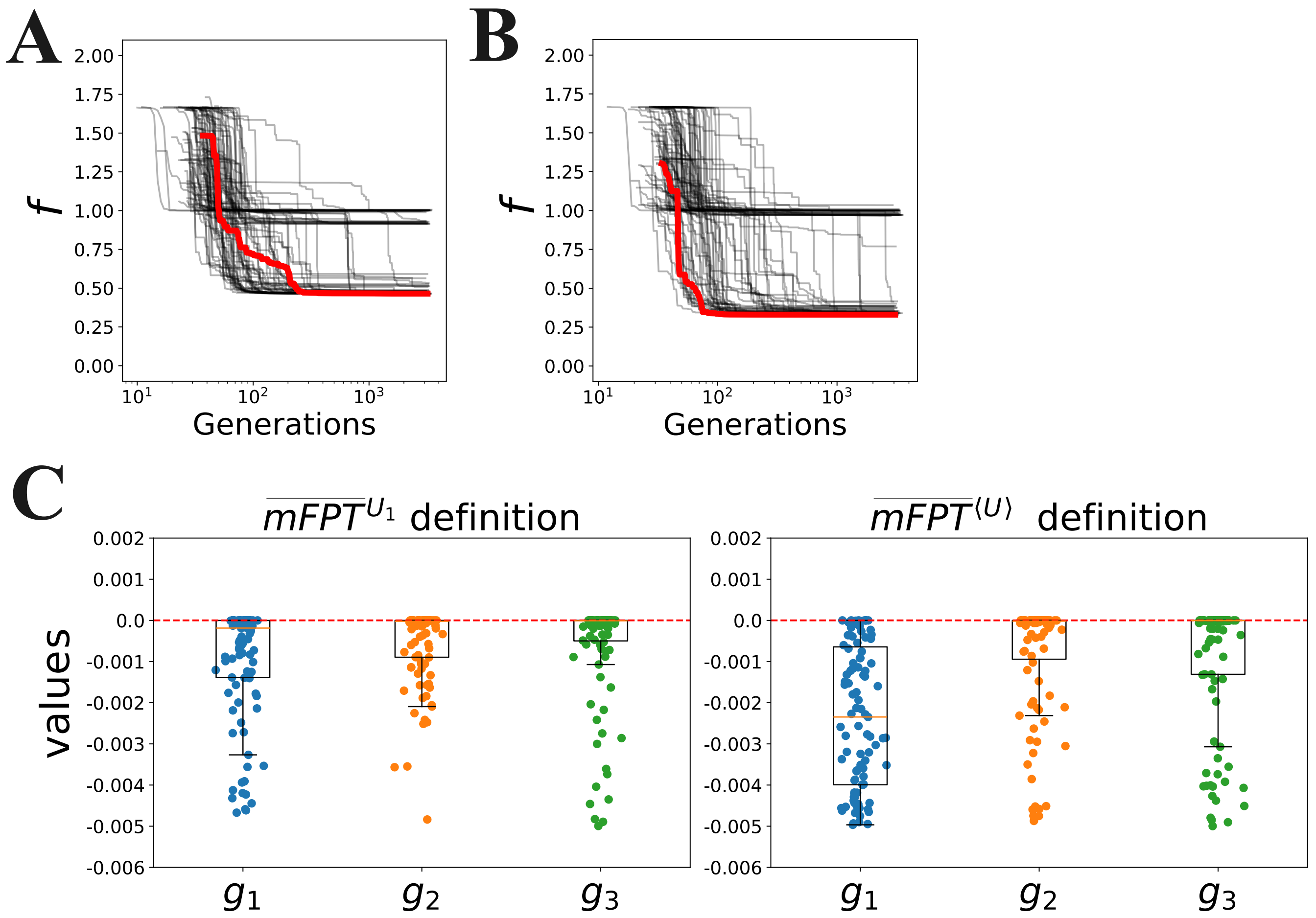


**Fig S2. Convergence of the optimization results in Fig. 3 and Fig. 4. (A)** Evolution of the coupling score, $f$, with the number of optimization generations for the parameter sets shown in Fig. 3 (with activation time defined as $\overline{\text{mFPT}}^{U_1}$). **(B)** Evolution of $f$ with the number of optimization generations for the parameter sets shown in Fig. 4 (with activation time defined as $\overline{\text{mFPT}}^{\langle U \rangle}$). In both panels, the red line represents the trajectory of the optimal parameter set with the least $f$. **(C)** Values of the constraint functions $g_1$, $g_2$ and $g_3$ at the end of each optimization. The $i$-th constraint is given by $g_i < 0$.

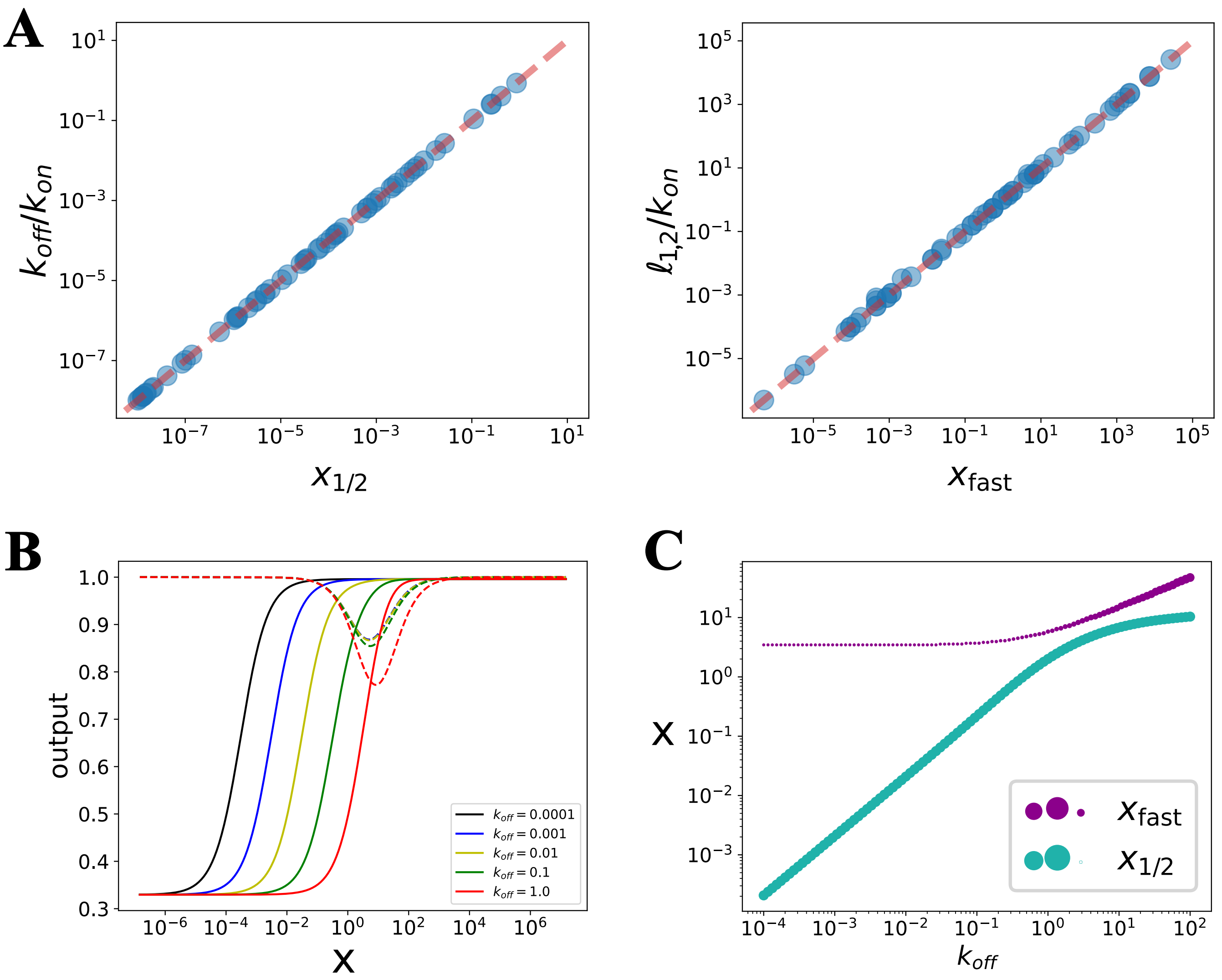


**Fig S3. Discrepancy between the concentration ranges over which $\overline{\mathrm{SS}}(x)$ and $\overline{\mathrm{mFPT}}^{U_1}(x)$ change significantly, for the parameter sets in Fig. 3D**. **(A)** *Left:* Comparison of $k_{\mathrm{off}}/k_{\mathrm{on}}$ against the concentration, $x_{1/2}$, at which $\overline{\mathrm{SS}}(x)$ is half-maximal. *Right:* Comparison of $\ell_{1,2}/k_{\mathrm{on}}$ against the concentration, $x_{\mathrm{fast}}$, at which $\overline{\mathrm{mFPT}}^{U_1}(x)$ is minimized. The red lines represent the loci at which each pair of quantities are equal. **(B)** $\overline{\mathrm{SS}}(x)$ and $\overline{\mathrm{mFPT}}^{U_1}(x)$ as functions of $x$ (solid lines and dashed lines, respectively), with each parameter set to the value in the best parameter set in Fig. 3D save for $k_{\mathrm{off}}$, which is varied as shown. **(C)** Values of $x_{1/2}$ and $x_{\mathrm{fast}}$ for the best parameter set in Fig. 3D, but with different values of $k_{\mathrm{off}}$. The size of each dot represents the value of $\Delta_{\overline{\mathrm{SS}}}$ (green) and $\Delta_{\overline{\mathrm{mFPT}}^{U_1}}$ (purple).

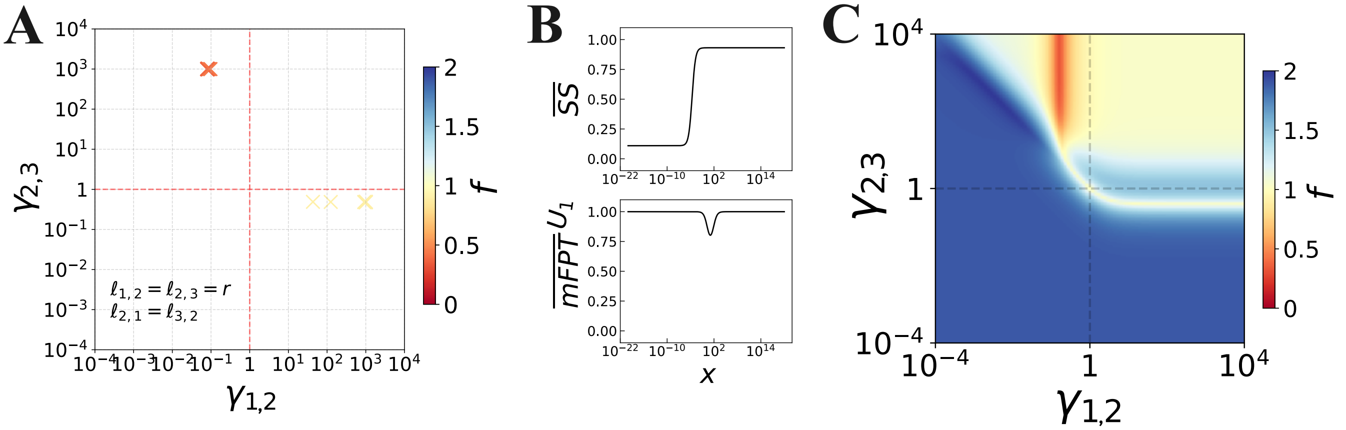


**Fig S4. Optimization with $\ell_{1,2} = \ell_{2,3} = r$ and $\ell_{2,1} = \ell_{3,2}$**, with $\gamma_{1,2}$ and $\gamma_{2,3}$ regulated. **(A)** Optimization results with $f < 1$ coloured by the value of $f$. **(B)** Responses for the best-isolated parameter set: $\ell_{1,2} = \ell_{2,3} = r = 9.999\delta$; $\ell_{2,1} = \ell_{3,2} = 23.93\delta$; $k_{on} = 0.403\delta/(1\text{c.u.})$; $k_{off} = 0.0001\delta$. **(C)** Scanning of the regulatory space for the best parameter set. The red area represents the minimum of the coupling score $f$.

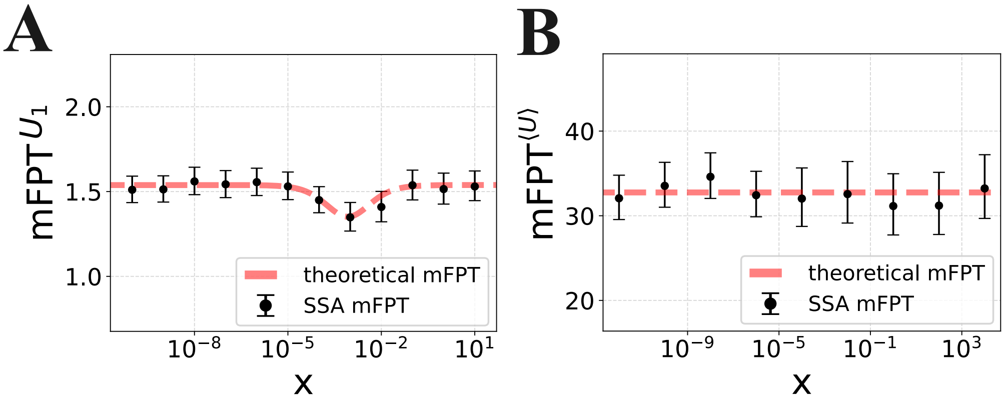


**Fig S5. Comparison between the theoretical and simulated activation times for the best parameter sets in Fig. 3-4.** **(A)**The simulated mFPT was obtained by simulating the stochastic process using the SSA algorithm in Gillespy2 [16]. The system is initialized in $U_1$, and simulated for a maximum simulation time approximately 100 times larger than the slowest horizontal rate in Fig. 3D, for 100 trajectories. For each of them, the time required to reach state $M$ is retrieved, and the average taken to yield $mFPT^{U_1}$. The error bar represents a 99% confidence interval of the mean. The theoretical value was obtained from Eqn 12. **(B)** Same as in A, but defining the activation time as $mFPT^{\langle U \rangle}$ and taking the best parameter set in Fig. 4E. We run an initial chain with initialization in $U_1$, waited approximately 100 times the slowest horizontal rate in Fig. 4E and then used the state of the chain to initialize another process with maximum time given by approximately 100 times the slowest horizontal rate in Fig. 4E, on which we computed the mean first passage time. The theoretical value was obtained from Eqn. 26 using Eqn. 12.

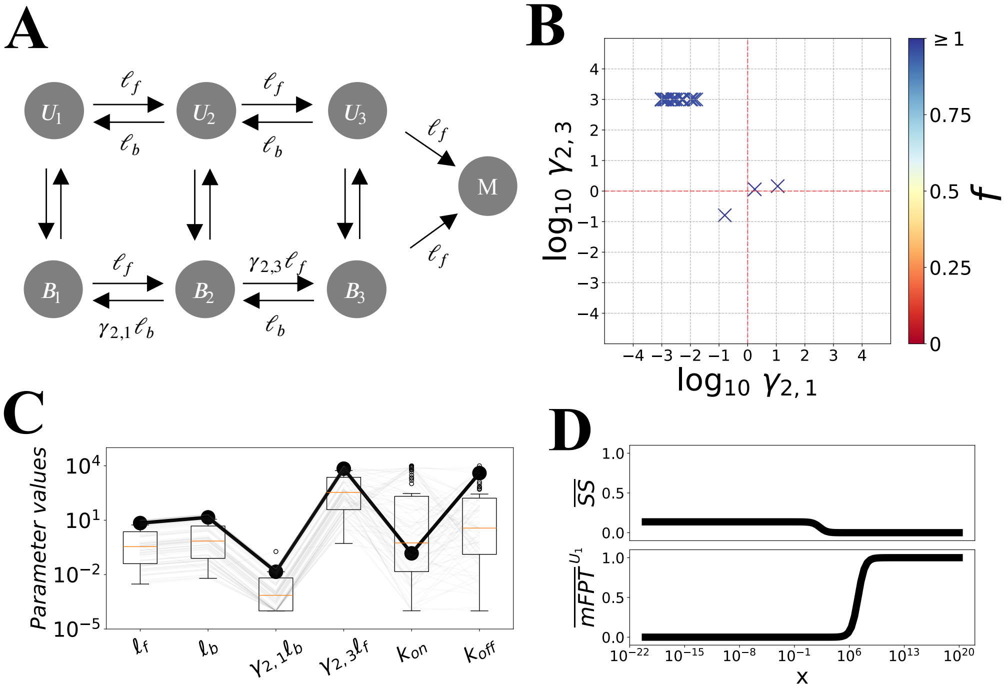


**Fig S6. Optimization results for $\ell_{1,2} = \ell_{2,3} = r$ and $\ell_{2,1} = \ell_{3,2}$**, with $\gamma_{2,1}$ and $\gamma_{2,3}$ regulated. **(A)** The optimization is performed imposing $\ell_{1,2} = \ell_{2,3} = a$ and $\ell_{2,1} = \ell_{3,2} = b$. **(B)** Within 100 optimization rounds, no parameter sets are found with a decoupling score significantly smaller than 1. **(C)** All the parameter sets are reported, with the black line denoting the best parameter set. Note that all entries are in units of $\delta$, except for $k_{\mathrm{on}}$ which is given in units of $\delta/(1\mathrm{c.u.})$. **(D)** Responses for the best parameter set ($\mathrm{argmin}_p\, f(p)$) clearly showing no decoupling.

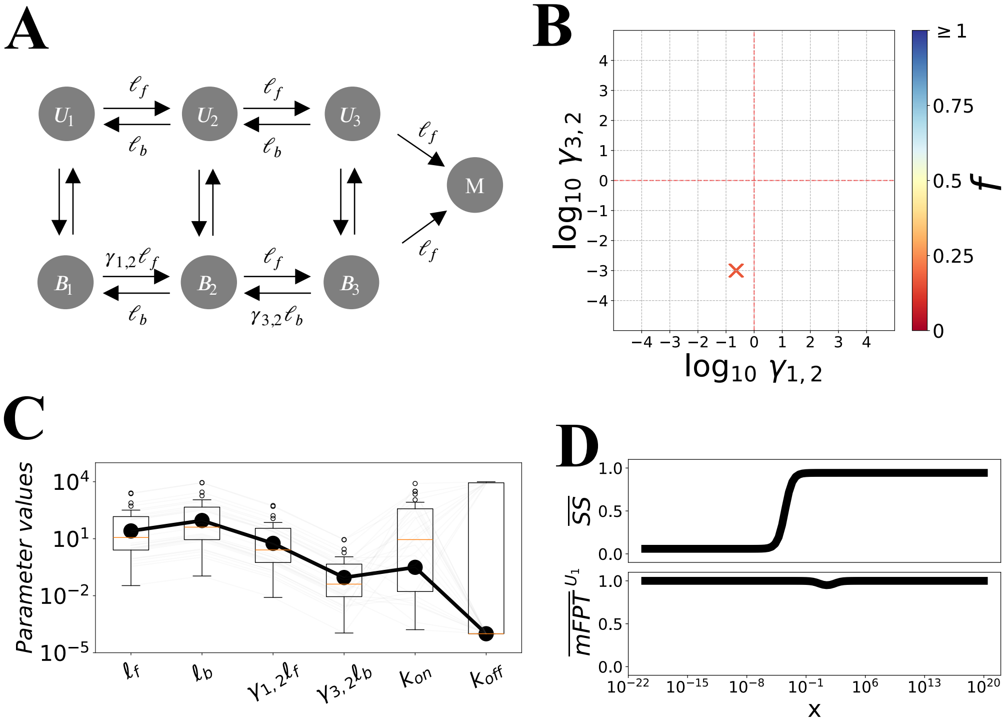


**Fig S7. Optimization results for $\ell_{1,2} = \ell_{2,3} = r$ and $\ell_{2,1} = \ell_{3,2}$,** with $\gamma_{1,2}$ and $\gamma_{3,2}$ regulated. **(A)-(D)**. Same as in the previous figure but assuming the effector is regulating the first forward transition ($B_1 \to B_2$) and the second backward transition ($B_3 \to B_2$). A clear decoupling is obtained in the incoherent regime, with $\gamma_{1,2} < 1$ and $\gamma_{3,2} < 1$, corresponding to a bias towards $B_1$ in the first transition and towards $B_3$ in the second (net deceleration of the first and net acceleration of the second transition).

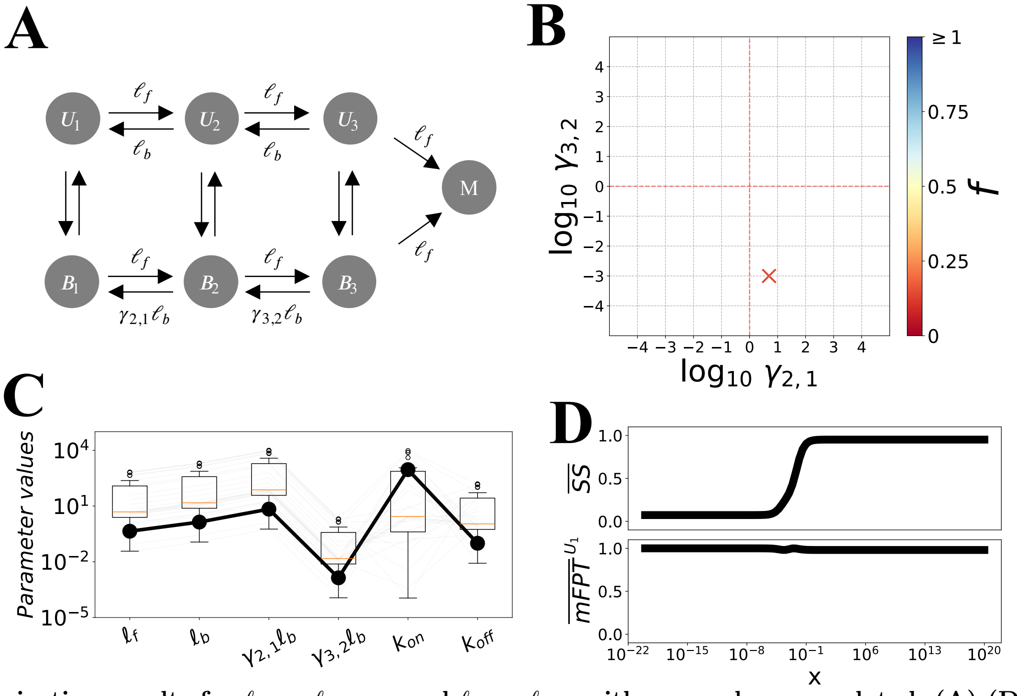


**Fig S8. Optimization results for $\ell_{1,2} = \ell_{2,3} = r$ and $\ell_{2,1} = \ell_{3,2}$, with $\gamma_{2,1}$ and $\gamma_{3,2}$ regulated**. **(A)-(D)**. Same as in the previous figure but assuming the effector is regulating the first backward transition ($B_2 \to B_1$) and the second backward transition ($B_3 \to B_2$). A clear decoupling is obtained in the incoherent regime, with $\gamma_{2,1} > 1$ and $\gamma_{3,2} < 1$, corresponding to a bias towards $B_1$ in the first transition and towards $B_3$ in the second (net deceleration of the first and net acceleration of the second transition).

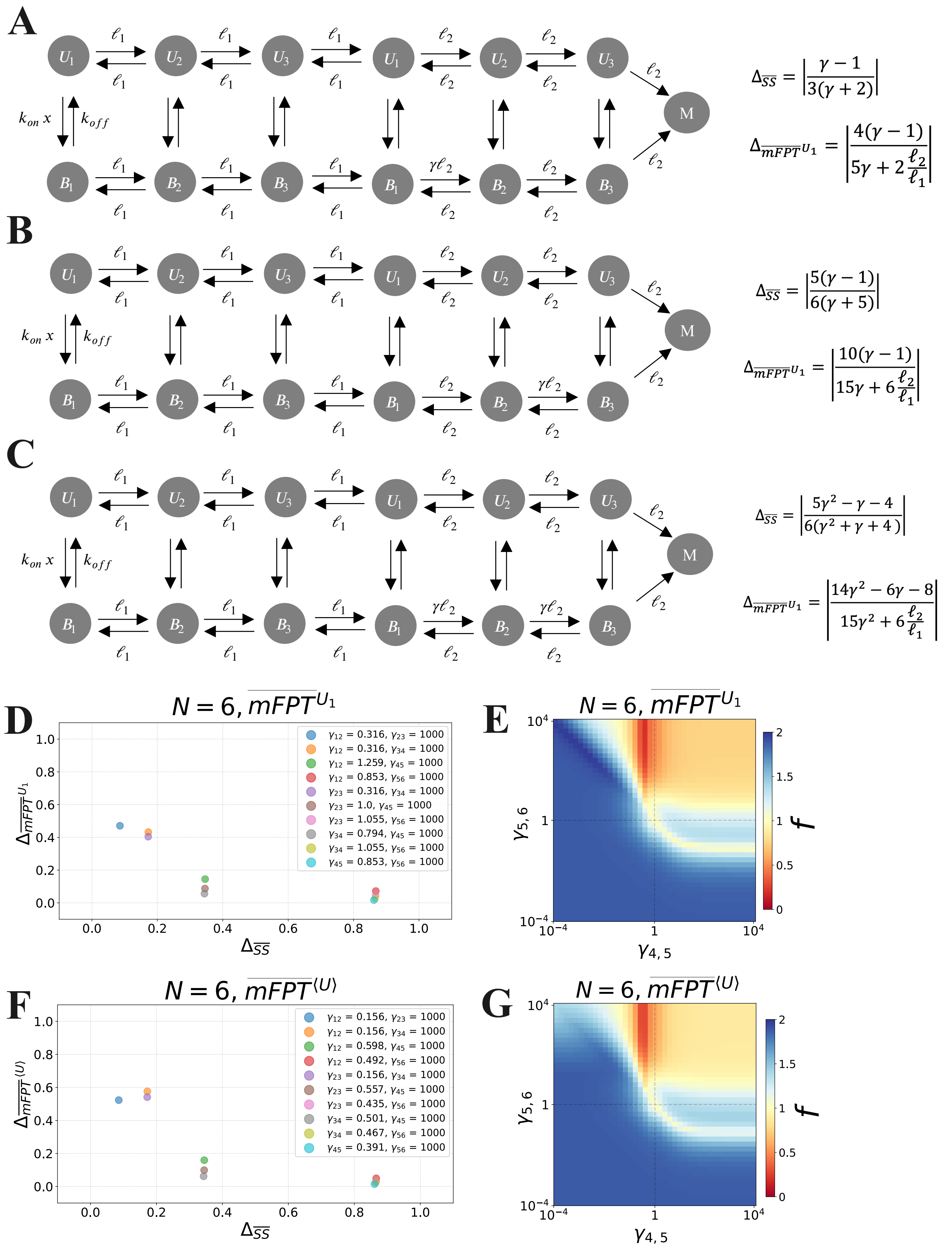


**Fig S9. Response decoupling for $N = 6$. (A)-(C)** Analytics for the parametrization in the graph on the left show that decoupling can be achieved under rate-scale separation ($\ell_2 >> \ell_1$) when regulation occurs on one or both of the fast transitions. *(continued on next page)*.

***(from previous page)*** **(D)** We extended the best parameter set obtained from the optimization under $N = 3$ and $RSC = 0.005$ from Fig. 3D. To extend the parameter set to $N = 6$ we started by computing the average rate between the forward transitions $\langle forward \rangle = (\ell_{1,2} + \ell_{2,3})/2$ and the backward transitions $\langle backward \rangle = (\ell_{2,1} + \ell_{3,2})/2$ and set the forward transitions in the $N = 6$ model (including $r$) equal to $\langle forward \rangle$, i.e. $\ell_{1,2}, \ldots, \ell_{5,6}, r = \langle forward \rangle$. Analogously we set all the backward transitions in $N = 6$ equal to the average backward transitions, i.e. $\ell_{2,1}, \ldots, \ell_{6,5} = \langle backward \rangle$. We preserved the rest of the original parameter set ($k_{on}$ and $k_{off}$). We then performed a first coarse-grained scanning of the regulatory space for each combination of two regulatory factors, followed by a local search to refine the regulatory values. The scatter plot shows the values of $\Delta_{\overline{SS}}$ and $\Delta_{\overline{mFPT}}$ for the best-identified solutions. Decoupling increases to the lower-right. **(E)** Regulatory parameter scanning for the regulatory factors showing the highest decoupling (highest $\Delta_{\overline{SS}}$ and lowest $\Delta_{\overline{mFPT}}$) in D. Notice that the minimum band is in the incoherent regulatory region. **(F)-(G)** Same as in D-E but assuming the system starts from an equilibrated system, such that activation time is defined with $mFPT^{\langle U \rangle}$. The best parameter set from Fig. 4E was extended as in (D).

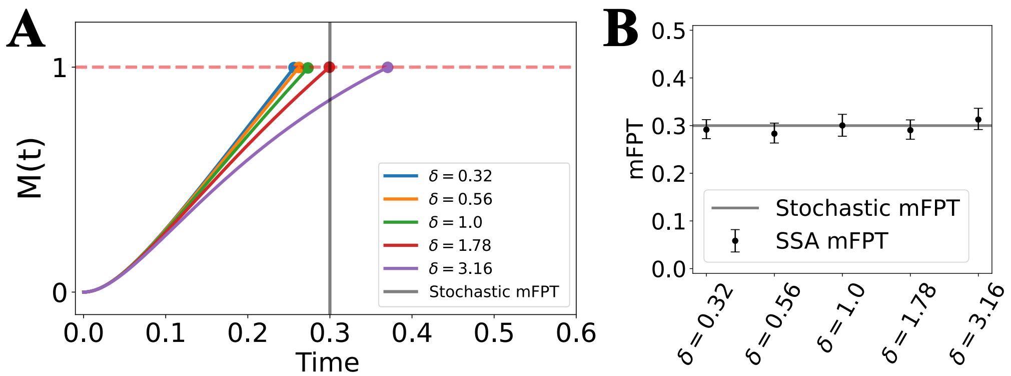


**Fig S10. Average time to produce one molecule (mFPT) vs time to produce on average one molecule. (A)**. Plot of Eq. S22 as a function of time, stopping when $M = 1$. The time required to reach $M = 1$ clearly depends on the degradation rate, with increasing $\delta$ corresponding to an increase in the activation time. On the contrary, the stochastic mFPT (from Eq. 12) does not depend on the degradation rate. The parameters used are: $a = b = r = 10$t.u. and the degradation rate according to the legend. **(B)**. Mean First Passage Time obtained from Gillespie SSA of the telegraph model. Error bars correspond to 99% confidence interval around the mean. The simulation time is 3 (t.u.) with $dt = 10^{-5}$ (t.u.), parameters as in $A$. The statistics is computed over 500 realizations.

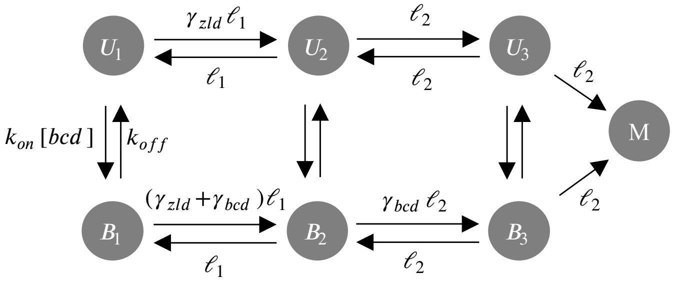


**Fig S11. Adaptation of the model $\mathcal{C}_3$ to account for the data in [11].** Two sets of ratescales are assumed, $\ell_1$ and $\ell_2$, and we assume rate scale separation with $\ell_2 \gg \ell_1$. We assume that the pioneer factor Zelda, not modelled explicitly given its constant concentration along the antero-posterior axis of the embryo, acts by increasing the magnitude of the slow transition $\ell_1$ by a positive factor $\gamma_{zld}$. Conversely, the morphogen Bicoid shows a concentration gradient along the anterior-posterior axis as recapitulated by the explicit Bicoid concentration dependence. Once Bicoid is bound, it modulates the transitions $B_i$ via the positive regulatory factor $\gamma_{bcd}$.

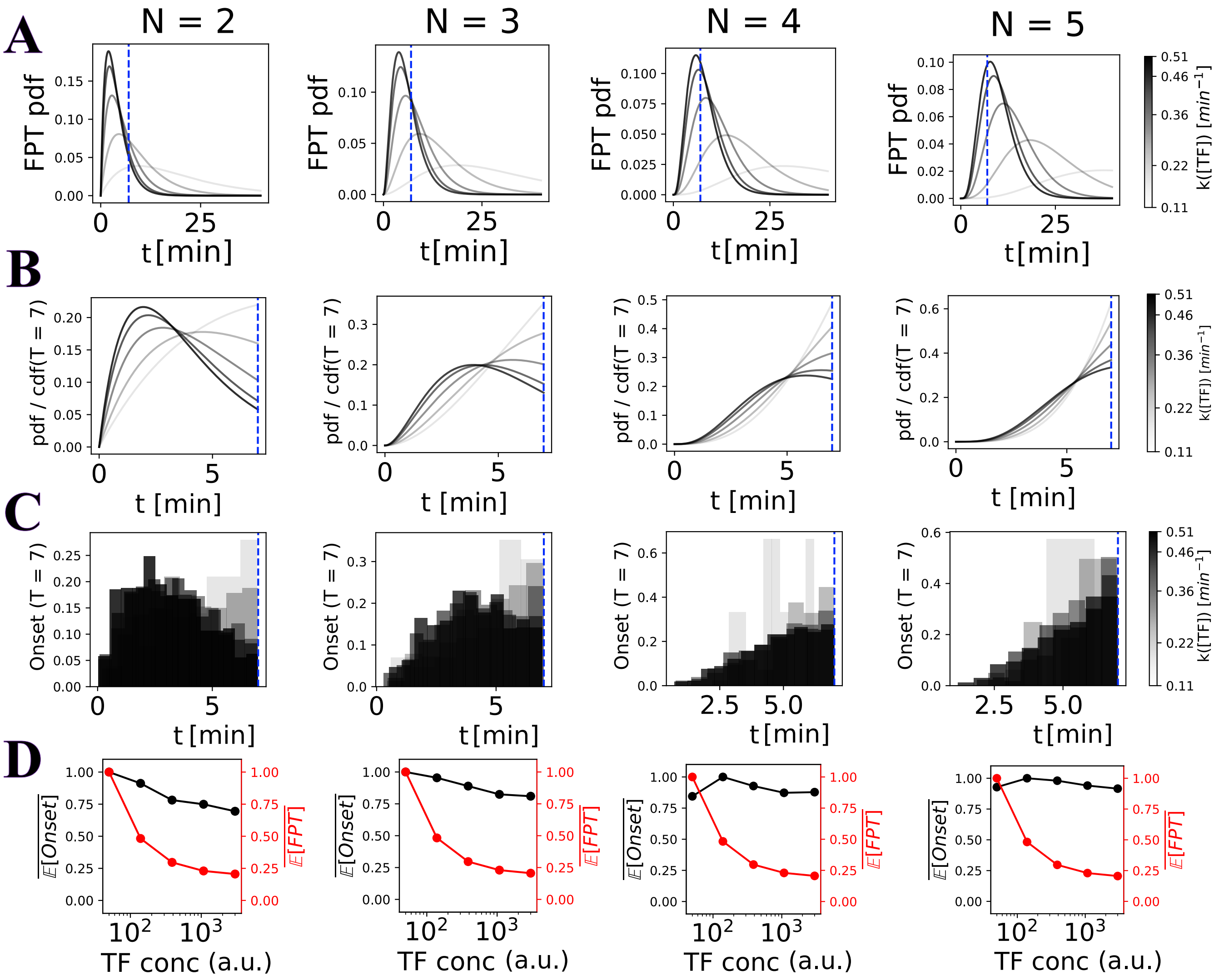


**Fig S12.** **(A)** First Passage Time distribution for the Erlang process with total number of states, $N + 1$. The rates are computed according to Supporting Information S7 with $c = 0.55$ $[min^{-1}]$ and $K_d = 250$ $[a.u.]$ as in [13]. The vertical line represents the length of the transcriptional active window ($t = 7$). **(B)** FPT distribution normalized by the value of the cumulative distribution at $t = 7$. **(C)** Gillespie SSA of the Erlang process with the parameters described in **A**. The onset time distribution of the traces that reach state $N$ within the simulation time ($t = 7$ $[min]$) is shown. **(D)** Average onset time and average FPT (considering the complete, untruncated distribution) normalized by their maximum as a function TF concentration as in Fig. 6 of [13]. The average onset time is computed from the distributions shown in **C**. The average first passage time is computed from Eqn. 12.
